# Functional interplay between condensin I and topoisomerase IIα in single-molecule DNA compaction

**DOI:** 10.1101/2025.01.21.634015

**Authors:** Yuko Tsubota, Keishi Shintomi, Kazuhisa Kinoshita, Yuki Masahara-Negishi, Yuuki Aizawa, Masami Shima, Tatsuya Hirano, Tomoko Nishiyama

**Affiliations:** Genome Integrity and Control Laboratory, Division of Biology, Graduate School of Science, Kyoto University, Sakyo, Kyoto, Japan; Chromosome Dynamics Laboratory, RIKEN, Wako, Saitama, Japan; Neural Circuit of Multisensory Integration RIKEN Hakubi Research Team, RIKEN Center for Brain Science, Wako, Saitama, Japan

## Abstract

Condensin I and topoisomerase IIα (topo IIα) are chromosomal ATPases essential for mitotic chromosome assembly. Mechanistically how the two ATPases cooperate to assemble mitotic chromosomes remains unknown. Here we use total internal reflection fluorescence microscopy to analyze the interplay between condensin I and topo IIα at single-molecule resolution. As observed in previous studies, condensin I alone predominantly forms DNA loops in an ATP-dependent manner. However, when topo IIα is included in the reaction, condensin I forms stable compact structures (termed “lumps”) in a manner dependent on the C-terminal domain of topo IIα. Each of the stable lumps contains a single condensin I complex and a single topo IIα dimer. Remarkably, we find that topo IIα, when catalytically active, renders the lumps resistant to protease treatment. Several lines of evidence show that the protease-resistant lumps contain knotted DNA. A mutant condensin I complex defective in ATP hydrolysis, together with topo IIα, forms smaller lumps in which the probability of DNA knotting is greatly reduced. Our results demonstrate how topo IIα-mediated strand passage is coupled with condensin I-mediated loop extrusion to generate a compact DNA structure. Together with recent studies, we discuss the functional implications of these observations in mitotic chromosome assembly and stabilization.

## Introduction

Mitotic chromosome assembly is a fundamental cellular process that ensures the faithful segregation of genetic information during the mitotic cell cycle. Extensive studies over the past decades led to a consensus that two distinct classes of ATPases, condensins and topoisomerase IIα (topo IIα), play central roles in this process^1,2^. For example, the chromosome scaffold, originally characterized as a proteinaceous structure observed in histone-depleted metaphase chromosomes, was later found to contain topo IIα^3,4^ and the subunits of condensin I^5,6^. More recent studies have successfully reconstituted the core reaction of this process *in vitro* by mixing a simple substrate with only six purified proteins, including core histones, three histone chaperones, topo IIα and condensin I^7,8^.

Eukaryotic topo IIα, which belongs to type IIA topoisomerase, introduces a double-strand break in one DNA strand and passes through a second DNA strand before rejoining the break in an ATP hydrolysis-dependent manner^9–11^. This strand passage reaction enables intermolecular catenation/decatenation and intramolecular knotting/unknotting of circular DNA *in vitro* (Fig. 1a). The direction of the reactions (i.e., catenation vs. decatenation; knotting vs. unknotting) depends on the topological states of the DNA substrates and the reaction environment. Such enzymology of topo IIα is reasonably well understood.

**Fig. 1.**
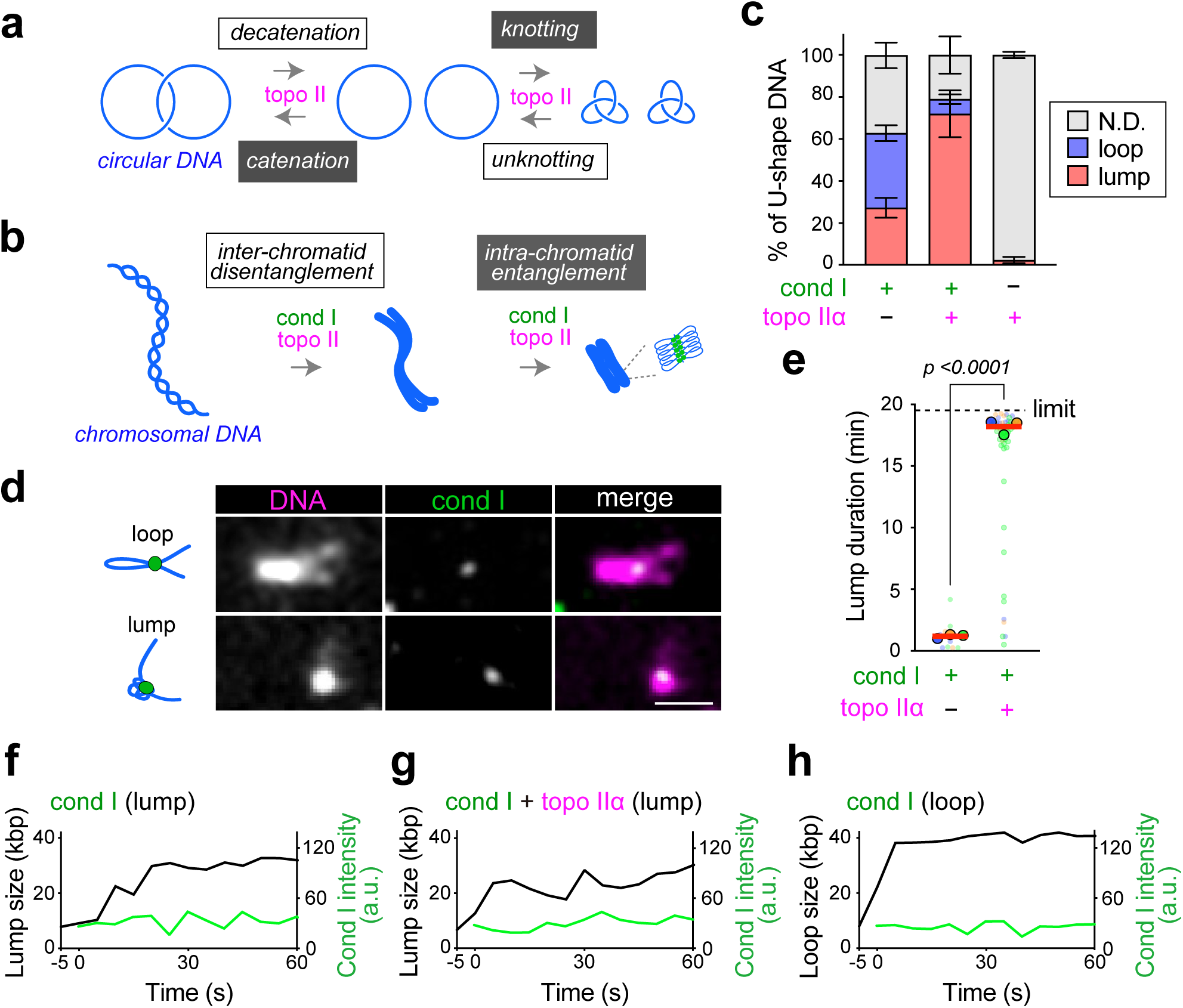
Condensin I forms stable lumps in the presence of topoisomerase IIα. **a**, The strand passage activity of topo IIα catalyzes the decatenation of catenated circular DNA or the catenation of free circular DNA. Alternatively, the same activity catalyzes the knotting of free circular DNA or the unknotting of knotted DNA. The black text/white boxes indicate topological simplifications (decatenation and unknotting), whereas the white text/black boxes indicate topological complications (catenation and knotting). **b**, At an early stage of chromosome assembly, the strand passage activity of topo IIα catalyzes inter-chromatid disentanglement, thereby promoting chromosome individualization and sister chromatid resolution. At a later stage of chromosome assembly, the same activity catalyzes intra-chromatid entanglement, making each chromatid more compact. The black text/white box indicates a topological simplification (inter-chromatid disentanglement), whereas the white text/black box indicates a topological complication (intra-chromatid entanglement). **c**, Frequency of lumps and loops on U-shape DNA in the presence of condensin I (cond I) alone, cond I and topo IIα, or topo IIα alone (cond I alone: n = 17, 28, 31 DNAs; cond I and topo IIα: n = 13, 12, 10 DNAs; topo IIα alone: n = 53, 79, 65 DNAs). Percentages of DNAs per condition (mean ± SEM) are plotted. Proteins were simultaneously introduced into U-shape DNA-tethered flow cells with imaging buffer at 20 μl/min flow for 100 sec. Unbound proteins were washed out using another imaging buffer flow during the observation. N.D. denotes that no significant structure was identified. **d**, Snapshots of loop and lump on a single lambda DNA. DNA was strained with SYTOX Orange and condensin I (cond I) was labeled by Halo-Alexa 488. Bar: 2 μm. **e**, Stability of lumps in the presence or absence of condensin I (cond I) and topo IIα. The duration of lump was measured under imaging buffer flow. The maximum duration is the observation time 20 min. The solid point denotes the median of each condition, and the red bar represents the mean of the median. Data were collected from 3 independent trials cond I alone: n = 4, 4, 6 DNAs; cond I and topo IIα: n = 13, 13, 16 DNAs). The P values were assessed by a two-tailed Mann-Whitney U test. **f-h**, Single-molecule tracking of condensin I (cond I) and compacted lump (f, g) or extruded loop (h) in the presence (g) or absence (f, h) of topo IIα and ATP. Time indicates that after cond I binding.

Much less is known about the mechanisms of action of condensin I. Early studies showed that condensin I has the ability to introduce positive superhelical tension into double-stranded DNA (dsDNA) *in vitro* in an ATP hydrolysis-dependent manner: relaxed circular DNA is converted into positively supercoiled DNA in the presence of type I topoisomerases^12^, whereas nicked circular DNA is converted into knotted DNA in the presence of a type II topoisomerase^13^. Because Cdk1 phosphorylation of condensin I strongly stimulates these activities^13,14^, they are thought to be physiologically relevant activities underlying mitotic chromosome assembly. Recent single-molecule studies have shown that condensins have the so-called loop extrusion activity that forms and expands DNA loops in an ATP hydrolysis-dependent manner^15,16^. The loop extrusion activity has been proposed to be a fundamental activity that organizes mitotic chromosomes by generating consecutive loops^17^.

As a natural consequence of the double-stranded nature of DNA, sister DNAs become entangled after DNA replication (Fig. 1b, left)^18^. Substantial evidence has demonstrated that the action of condensins helps topo II to facilitate the resolution of such entanglements (inter-chromatid disentanglement) in preparation for complete sister chromatid separation during mitosis (Fig. 1b, center)^19,20^. Interestingly, evidence is now accumulating for a second function of topo IIα in mitotic chromosome assembly: DNA strands within individualized chromatids become self-entangled (intra-chromatid entanglement) through the action of topo IIα (Fig. 1b, right)^8,21,22^. It is speculated that this previously underappreciated action of topo IIα is facilitated by the crowded environment created during the process of mitotic chromosome assembly^8^.

Although the individual activities of topo IIα and condensin I have been studied extensively, only a limited number of efforts have been made to address the question of how the two ATPases cooperate to assemble mitotic chromosomes at a mechanistic level^23^. In the current study, we have used total internal reflection fluorescence microscopy to analyze the interplay between condensin I and topo IIα at single-molecule resolution. To this end, topo IIα is added to a standard condensin I-mediated loop extrusion assay. We find that inclusion of topo IIα converts the condensin I-mediated DNA structures into stable compact structures in a manner that is dependent on the C-terminal domain (CTD) of topo IIα. Moreover, the strand passage activity of topo IIα induces DNA knotting within the stable lumps, rendering them resistant to protease treatment. ATP hydrolysis by condensin I stimulates the topo IIα-mediated knotting reaction. Our results demonstrate how topo IIα-mediated strand passage is coupled with condensin I-mediated loop extrusion to generate a compact DNA structure. The functional implications of these observations for mitotic chromosome assembly and stabilization are discussed.

## Results

### Condensin I forms stable lumps in the presence of topo IIα

To address the question of how condensin I and topo IIα cooperate to change the conformation of DNA at a single molecule resolution, we introduced topo IIα to our condensin I-mediated loop extrusion assay. In brief, we utilized a flow cell-based assay system, which was visualized under a total internal reflection fluorescence microscope (TIRFM) as previously reported^24^. 48.5-kbp λDNA was biotinylated at both ends and tethered onto a streptavidin-coated coverslip to make U-shape DNA substrates. A mammalian condensin I complex and *Xenopus laevis* topo IIα were expressed by using the baculovirus expression system and purified as previously reported^8,25^. The purified condensin I was then fluorescently labeled.

When condensin I alone (1 nM) was injected into the flow cells, U-shape DNA was either converted into a loop (35.6%) as reported previously^15,16,25,26^ or a compact structure that consisted of a DNA mass without extruded loop (27.3%), hereafter referred to as a “lump” (Fig. 1c, see cartoon in Fig. 1d). Injection of topo IIα alone at a low concentration (0.125 nM) did not change the DNA shape. However, when condensin I and topo IIα were injected together at concentrations of 1 nM and 0.125 nM, respectively, lump formation dominated: lumps were formed on ∼70% of U-shape DNA (see an example in Fig. 1d, “lump”), whereas loops were formed on less than 10% of DNA (see an example in Fig. 1d, “loop”). Importantly, we found that the lumps formed by condensin I and topo IIα were sustained for far longer time (in most cases, sustained throughout the observation time [20 min]) than those formed by condensin I alone (Fig. 1e), although the kinetics of lump formation were similar under the two conditions (Figs. 1f-g). Time-lapse imaging revealed that DNA began to be converted into lumps immediately after condensin I binding (Figs. 1f, and Extended Data Figs. 1a-d, Supplemental movie S1), and the lumps grew more slowly than the loops (Figs. 1f-h). We did not observe examples of loop-to-lump conversion unless a second condensin is bound to the loops (Supplemental movie S2), suggesting that the lump formation is initiated before loop growth.

In the absence of ATP, we observed only limited numbers of lumps with reduced size (Figs. 2a-b, and Extended Data Figs. S2a-b) and reduced rates (Fig. 2c), indicating that the stable lump formation is ATP-dependent.

**Fig. 2.**
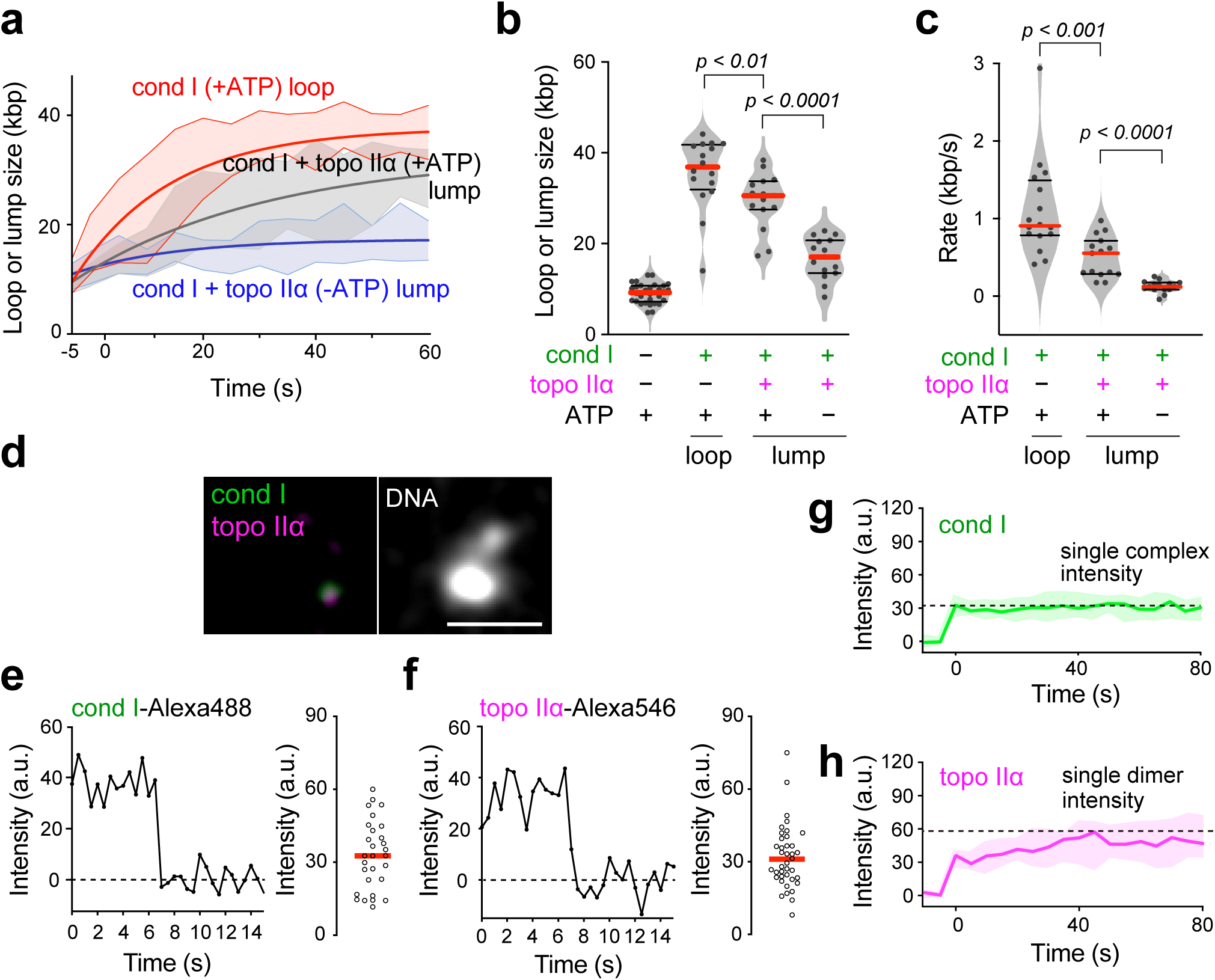
Tracking of single DNA behaviors in the presence of condensin I and topo IIα. **a**, Time trace of loop or lump size associated with single condensin I (cond I) complexes in the presence or absence of topo IIα. Exponential fittings with interquartile ranges are shown. Data were collected from 15 samples in more than 2 independent trials. **b**, Size of lump or loop formed by condensin I (cond I) in the presence or absence of topo IIα and ATP. The red and black bars denote the median and interquartile, respectively. n = 45, 15, 15, 15 DNAs analyzed (from left to right). The P values were assessed by two-tailed Mann-Whitney U test. **c**, Rate of loop or lump formation by condensin I (cond I) in the presence or absence of topo IIα and ATP. The red and black bars denote the median and quartiles, respectively. n = 15, 15, 15 DNAs analyzed (from left to right). The P values were assessed by a two-tailed Mann-Whitney U test. **d**, A snapshot of condensin I (cond I)^A488^ and topo IIα^A546^ on a lump. Cond I^A488^ (1 nM) and topo IIα^A546^ (50 pM) were injected simultaneously and washed out with imaging buffer (IB) after 100 sec of injection. After the detection of cond I^A488^, DNA was strained with SYTOX Green. Bar: 2 μm. **e-f**, Time trace of cond I^A488^ (e) and topo IIα^A546^ (f) signals binding non-specifically to the glass surface. Cond I^A488^ or topo IIα^A546^ particles were bleached by 488 nm or 561 nm laser for over 15 sec and the fluorescent images were acquired every 0.5 sec. Note that both signals were bleached in one step (left). Fluorescence intensities of particles bleached in one step were plotted (right). **g-h**, Binding kinetics of cond I^A488^ (g) and topo IIα^A546^ (h) on DNA. Dashed lines denote fluorescence intensities of single cond I^A488^ complexes (g) or single topo IIα^A546^ dimers (h). Fluorescence intensities of cond I^A488^ and topo IIα^A546^ were measured every 5 sec for 150 sec. Time 0 indicates the timing when the first cond I^A488^ bound to DNA. Median ± interquartile ranges are shown. n = 11 (g), 16 (h) DNAs analyzed.

To identify how many molecules of condensin I and topo IIα are involved in the formation of stable lumps, we measured the intensities of the fluorescently labeled condensin I (Alexa488) and topo IIα (Alexa546) that bind to each lump. In most cases, single condensin I and single topo IIα particles were observed in each lump (Fig. 2d). By comparing the intensities of these single particles with those of single condensin I and single topo IIα molecules that were bleached in one step by photobleaching experiments (Figs. 2e-f), we found that each of the stable lumps contains a single condensin I complex and a single topo IIα dimer (Figs. 2g-h, Extended Data Fig. 2c).

### Condensin I/topo IIα-mediated lumps are resistant to protease treatment

Although the lumps formed by condensin I alone were far less stable than those formed by both condensin I and topo IIα (Fig. 1e), we found that condensin I was continued to be detectable on DNA throughout the observation time [20 min] even after the lumps were dissolved (Extended Data Fig. 3a). Thus, condensin I binds stably to DNA even in the absence of topo IIα.

To further clarify the differences between condensin I-mediated and condensin I/topo IIα-mediated lumps, we then treated the two types of lumps with a protease (Proteinase K). As expected, the fluorescent signals of both condensin I and topo IIα disappeared from DNA upon protease treatment (Extended Data Figs. 3b-c). We found that the lumps and loops formed by condensin I alone were immediately dissolved upon this treatment (Figs. 3a upper, and 3b-c). Remarkably, however, the lumps formed by both condensin I and topo IIα remained even after protease treatment although their size decreased slightly (Figs. 3a lower, and 3b-c). These lumps are hereafter referred to as “protease-resistant lumps”. We also found that the small, condensin I/topo IIα-mediated lumps observed in the absence of ATP (Fig. 2b) were protease-sensitive (Fig. 3c).

**Fig. 3.**
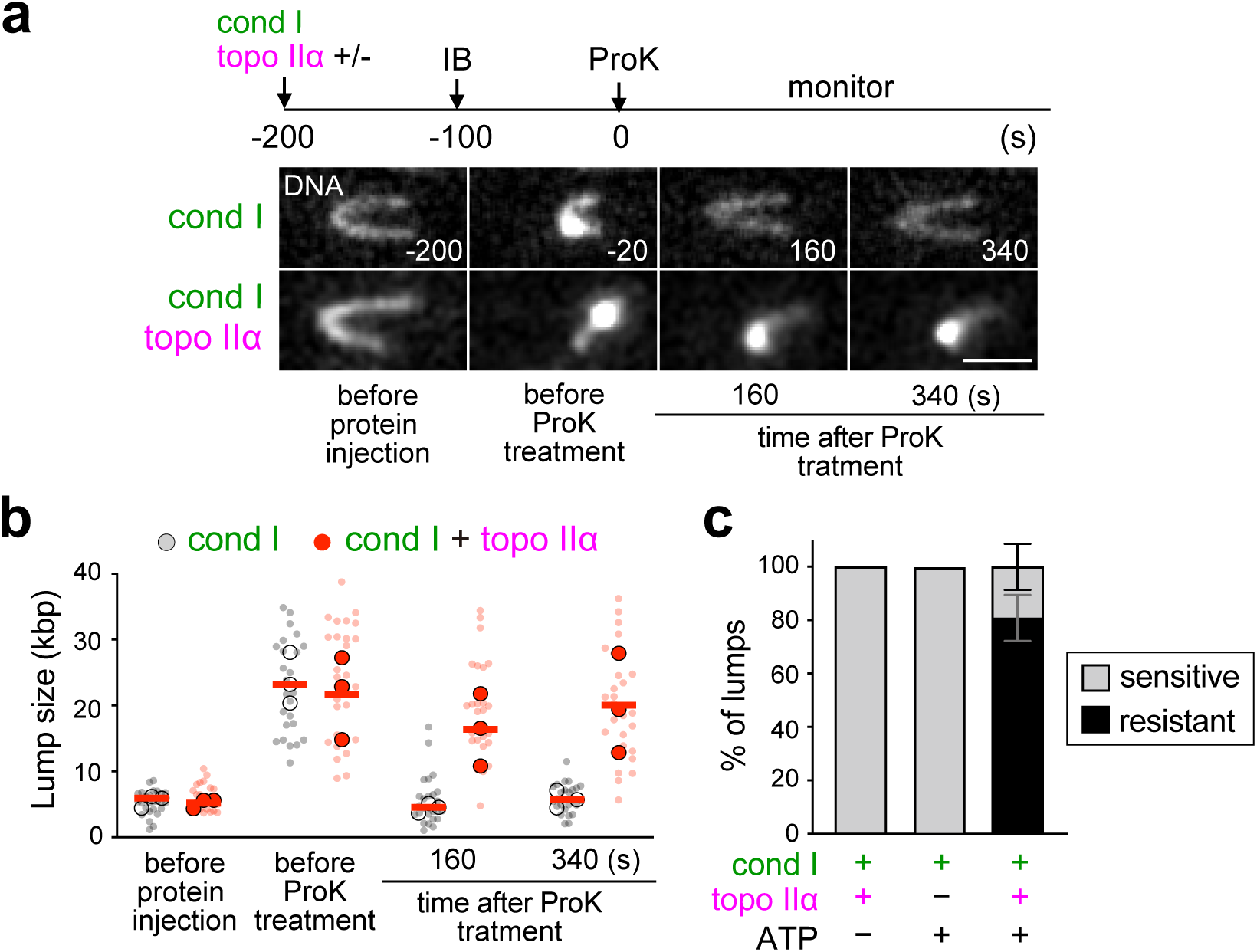
Condensin I/topo IIα-mediated lumps are resistant to protease treatment. **a**, Snapshots of the lumps formed by condensin I (cond I) alone or cond I and topo IIα before (−20 sec) and after (160 sec, 340 sec) Proteinase K (ProK) treatment. Cond I and topo IIα were injected simultaneously (−200 sec), washed out with buffer (IB) (−100 sec), and then ProK (0.5 mg/mL) was injected (0 sec). Times (sec) after ProK treatment are shown. **b**, The size of the lumps formed in the experiment shown in (a). The solid point denotes the median of each condition, and the bar denotes the mean of the median. Data were collected from 3 independent trials (condensin I (cond I) alone: n = 10, 9, 7 DNAs; cond I and topo IIα: n = 11, 10, 7 DNAs). **c**, Frequency of ProK-resistant and sensitive lumps associated with single condensin I (cond I) complexes in the presence or absence of ATP and topo IIα. Data were collected from 3 independent trials (cond I alone with ATP: n = 3, 14, 5 DNAs; cond I and topo IIα without ATP: n = 6, 4, 17 DNAs; cond I and topo IIα with ATP: n = 11, 10, 9 DNAs, mean ± SEM).

Although ∼80% of the lumps formed by condensin I and topo IIα were protease-resistant, ∼20% were judged to be protease-sensitive (Fig. 3c). We then compared the DNA binding kinetics of condensin I and topo IIα between the protease-resistant and protease-sensitive lumps, and found a tendency for the lumps to become more protease-resistant when condensin I bound to DNA before topo IIα (Extended Data Fig. 4a). These results suggested that condensin I creates a preferred DNA structure for topo IIα to bind and thereby generates the protease-resistant lumps. This idea was further supported by experiments involving the sequential addition of condensin I and topo IIα (Extended Data Fig. 4b).

Finally, we examined changes in the size of protease-resistant lumps (after deproteinization) in response to DNA tension imposed by different flow rates. If the DNA within the lumps was topologically constrained, then it was expected that 1) the lumps would be maintained even as DNA tension increased; and 2) the apparent lump size would decrease as DNA tension increased, whereas it would recover as tension decreased. We evaluated the apparent lump size by measuring the cross-sectional intensity of the lumps and comparing either the highest DNA intensity on the lump (Max) or the sum of the DNA intensities on the lump (Area Under the Curve [AUC]), and found that both Max and AUC values decreased when the flow rate was increased, but they were restored when the flow rate was decreased again (Extended Data Fig. 5). The lump size was approximately 2∼3-fold greater in compacted tips than that in uncompacted tips both before and after extension. These results strongly suggested that the condensin I/topo IIα-mediated lump, but not the condensin I-mediated lump, contains a topologically constrained DNA structure.

### DNA strand passage by topo IIα is essential for protease-resistant lump formation

We next directly tested whether the DNA strand passage activity of topo IIα is required for protease-resistant lump formation. We expressed and purified *Xenopus laevis* topo IIα harboring the strand passage-deficient mutation Y803F (Extended Data Fig. 6a)^27,28^. We confirmed that the mutant topo IIα (topo IIα^Y803F^) was catalytically null in the standard decatenation assay *in vitro* (Extended Data Fig. 6b). As expected, topo IIα^Y803F^ bound to chromatin but failed to assemble mitotic chromosomes in *Xenopus* egg extracts (Extended Data Fig. 6c).

Topo IIα^Y803F^ was then subjected to our single-molecule assay. We found that topo IIα^Y803F^ could bind to DNA (Extended Data Fig. 6d) and supported lump formation similarly to topo IIα^WT^ in terms of formation efficiency (Fig. 4a), kinetics (Fig. 4b), stability (Fig. 4c), size (Fig. 4d), and formation rate (Fig. 4e). These results are consistent with the observation that topo IIα^Y803F^ retains its ability to bind to chromatin in *Xenopus* egg extracts as described above. Remarkably, however, the lumps formed by topo IIα^Y803F^ were largely protease-sensitive, as opposed to those formed by topo IIα^WT^ (Fig. 4f). Thus, the strand passage activity is essential for protease-resistant lump formation, but not for stable lump formation. However, despite the fact that the topo IIα^Y803F^ is catalytically null, a non-negligible fraction (∼10%) of lumps were judged to be protease-resistant under the current condition (Fig. 4f). We reasoned that the DNA present in the residual protease-resistant lumps might be constrained in a non-topological manner. In fact, when a higher extension force was imposed on the DNA by increasing the flow rate (from 20 μl/min to 1 ml/min), the residual protease-resistant lumps formed by topo IIα^Y803F^ completely disappeared (Extended Data Fig. 6e, YF), although the protease-resistant lumps formed by topo IIα^WT^ were maintained (Extended Data Fig. 6e, WT).

**Fig. 4.**
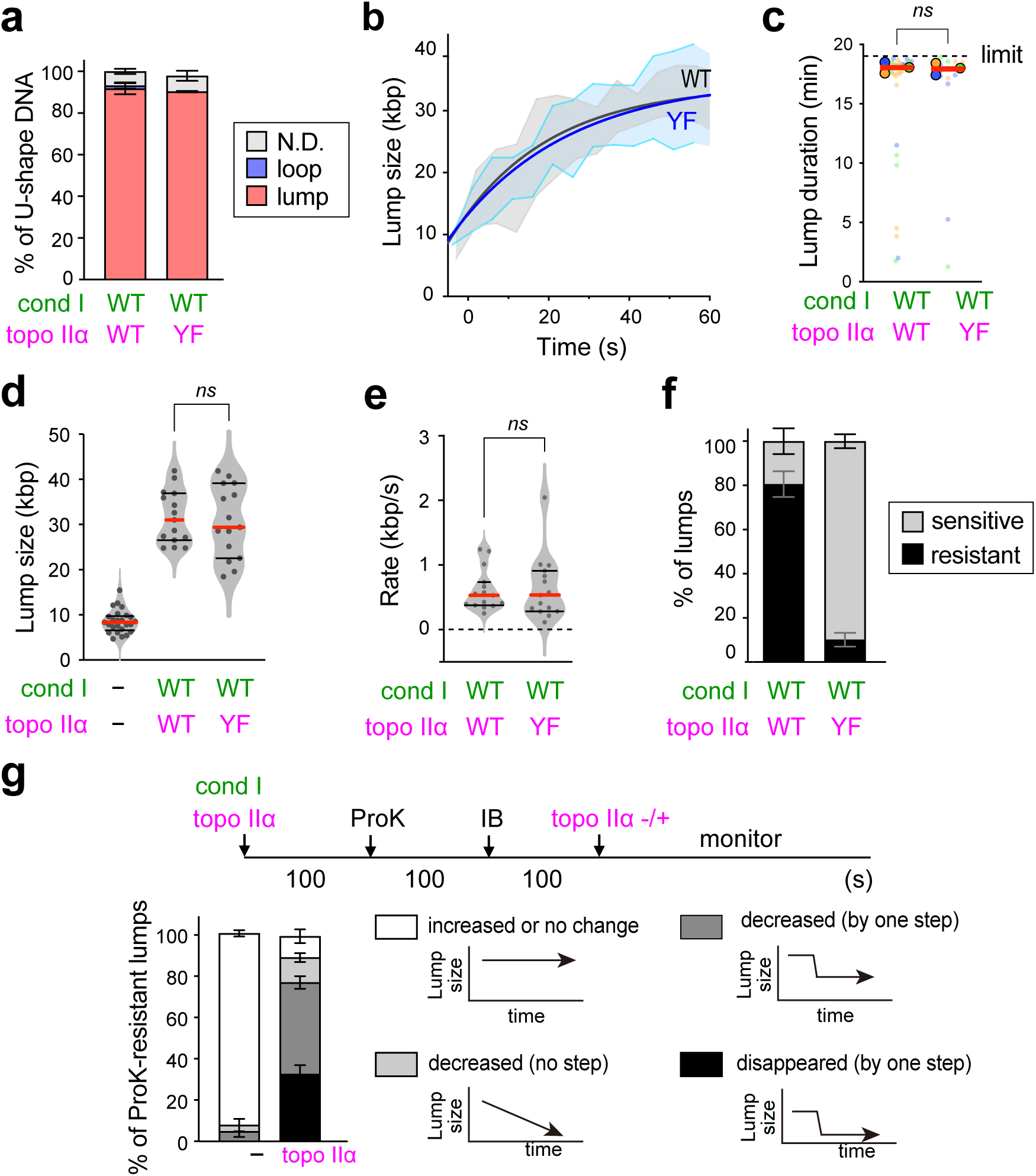
DNA strand passage by topo IIα is essential for protease-resistant lump formation. **a**, Frequency of lumps and loops on U-shape DNA formed by condensin I (cond I) and either topo IIα^WT^ (WT) or topo IIα^Y803F^ (YF). Data were collected from 3 independent trials (topo IIα^WT^: n = 22, 20, 16 DNAs; topo IIα^Y803F.^: n = 20, 11, 10 DNAs). **b**, Time trace of the size of a lump formed by condensin I in the presence of either topo IIα^WT^ (WT) or topo IIα^Y803F^ (YF). Exponential fitting lines with interquartile ranges are shown. Data were collected from 15 DNAs from 3 independent trials. **c**, Stability of lump in the presence of condensin I (cond I) and either topo IIα^WT^ (WT) or topo IIα^Y803F^ (YF). The duration of lump to be maintained was measured under imaging buffer flow. The maximum duration is the observation time 20 min. The solid point denotes the median of each condition, and the red bar represents the mean of the median. Data were collected from 3 independent trials (topo IIα^WT^: n = 10, 13, 9 DNAs; topo IIα^Y803F.^: n = 7, 4, 5 DNAs). The P values were assessed by a two-tailed Mann-Whitney U test. **d**, Size of lump formed in the presence or absence of condensin I (cond I) and either topo IIα^WT^ (WT) or topo IIα^Y803F^ (YF). The red and black bars denote the median and interquartile, respectively (n = 30, 15, 15 DNAs). The P values were assessed by a two-tailed Mann-Whitney U test. **e**, Rate of lump formation in the presence of condensin I (cond I) and either topo IIα^WT^ (WT) or topo IIα^Y803F^ (YF). The red and black bars denote the median and interquartile, respectively (n = 15 (topo IIα^WT^), 15 (topo IIα^Y803F^), The P values were assessed by a two-tailed Mann-Whitney U test). **f**, Frequency of protease-resistant and -sensitive lumps associated with single condensin I (cond I) complexes in the presence of either topo IIα^WT^ (WT) or topo IIα^Y803F^ (YF). Data were collected from 3 independent trials (topo IIα^WT^: n = 19, 19, 15 DNAs; topo IIα^Y803F.^: n = 18, 10, 9 DNAs; mean ± SEM). **g**, Changes in protease-resistant lumps after the 2nd topo IIα treatment. Condensin I (cond I) (1 nM) and topo IIα (0.125 nM) were injected simultaneously, the DNAs were treated with protease (ProK) for 100 sec, washed out with imaging buffer (IB). Then IB (-) or the 2nd fresh topo IIα (0.25 nM) was injected again and the lump sizes were monitored for 15 min. Changes in the lump size over time were categorized into 4 groups (white: increase or no change, right gray: gradual decrease, dark gray: decreased by one step, black: disappeared by one step). Data were collected from 3 independent trials with at least 19 lumps per condition (mean ± SEM).

We next treated deproteinized lumps with freshly added topo IIα and found that most of the lumps diminished or disappeared (Fig. 4g). It is noteworthy that in most cases, DNA intensities were decreased in a stepwise manner (Fig. 4g), implying that these lumps were dissolved by one or a few step(s) of the strand passage reaction. Based on these results, we concluded that the protease-resistant lumps generated by condensin I and topo IIα^WT^ contain DNA knots, as reported in the previous bulk biochemical studies^13,29^.

### The CTD of topo IIα is required for protease-resistant lump formation

To get additional insights into the mechanism of protease-resistant lump formation, we next asked if stable binding of topo IIα to DNA is required for this process. Previous studies have shown that the intrinsically disordered CTD of topo IIα is required for stable localization of topo IIα to reconstituted mitotic chromosomes and their thickening^8^. We prepared fluorescently labeled topo IIα lacking its CTD (topo IIα^ΔCTD^; Fig. 5a) and found that the addition of topo IIα^ΔCTD^ had little, if any, impact on condensin I-mediated structural changes of DNA in our single-molecule assay (Fig. 5b). Similar to the lumps formed with condensin I alone, the lumps formed with condensin I and topo IIα^ΔCTD^ were all protease-sensitive (Fig. 5c). Thus, stable DNA binding mediated by the CTD of topo IIα is required for protease-resistant lump formation although topo IIα^ΔCTD^ is catalytically fully active as judged by the standard DNA decatenation assay^8^.

**Fig. 5.**
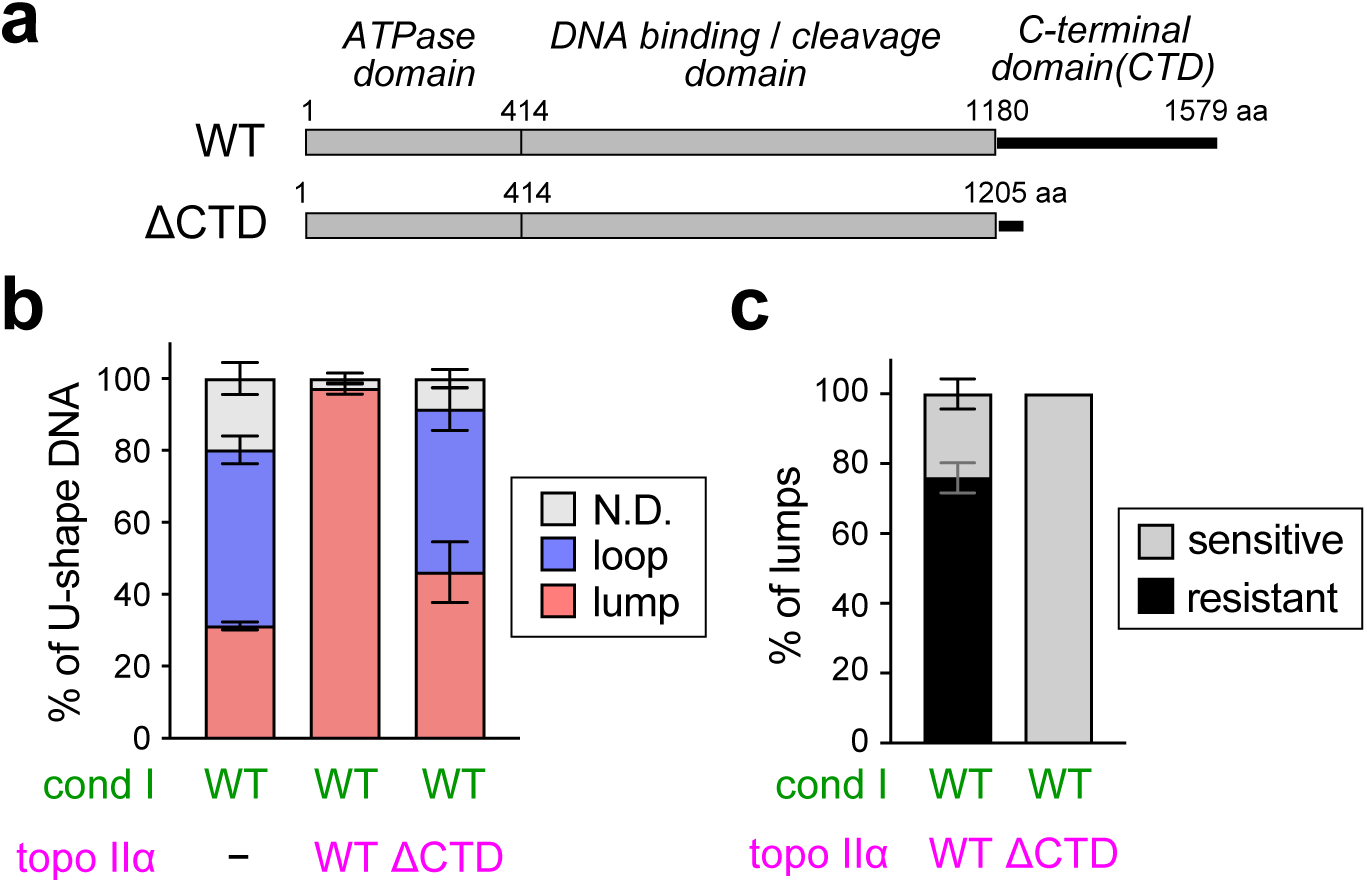
The C-terminal domain of topo IIα is required for protease-resistant lump formation. **a**, Schematic representation of topo IIα^WT^(WT) and topo IIα^ΔCTD^ (ΔCTD). **b**, Frequency of lumps and loops on U-shape DNA formed by condensin I (cond I) in the presence of either topo IIα^WT^(WT) and topo IIα^ΔCTD^ (ΔCTD). Data were collected from 3 independent trials (cond I alone: n = 67, 30, 46 DNAs; cond I and topo IIα^WT^: n = 32, 19, 17 DNAs; cond I and topo IIα^ΔCTD^: n = 23, 37, 38 DNAs, mean ± SEM). **c**, Frequency of protease-resistant and sensitive lump formation by condensin I (cond I) and either topo IIα^WT^(WT) and topo IIα^ΔCTD^ (ΔCTD). Data were collected from 3 independent trials topo IIα^WT^: n = 31, 18, 16 DNAs; topo IIα^ΔCTD^: n = 14, 17, 12 DNAs, mean ± SEM).

### ATP hydrolysis by condensin I strongly stimulates protease-resistant lump formation

Next we investigated how condensin I’s ATPase activity affects lump formation. For this purpose, we prepared a fluorescently labeled, condensin I mutant complex deficient in ATP hydrolysis (Smc2[E1114Q] and Smc4 [E1218Q], henceforth referred to as condensin I^EQ^)^25^. Consistent with previous studies in all types of eukaryotic SMC complexes^16,30–32^, condensin I^EQ^ alone showed no loop extrusion activity (Fig. 6a). Moreover, topo IIα-independent lump formation was also significantly suppressed (Fig. 6a). On the contrary, in the presence of topo IIα, condensin I^EQ^ formed lumps with a high frequency and stability indistinguishable from those observed with condensin I^WT^ (Figs. 6b-c).

**Fig. 6.**
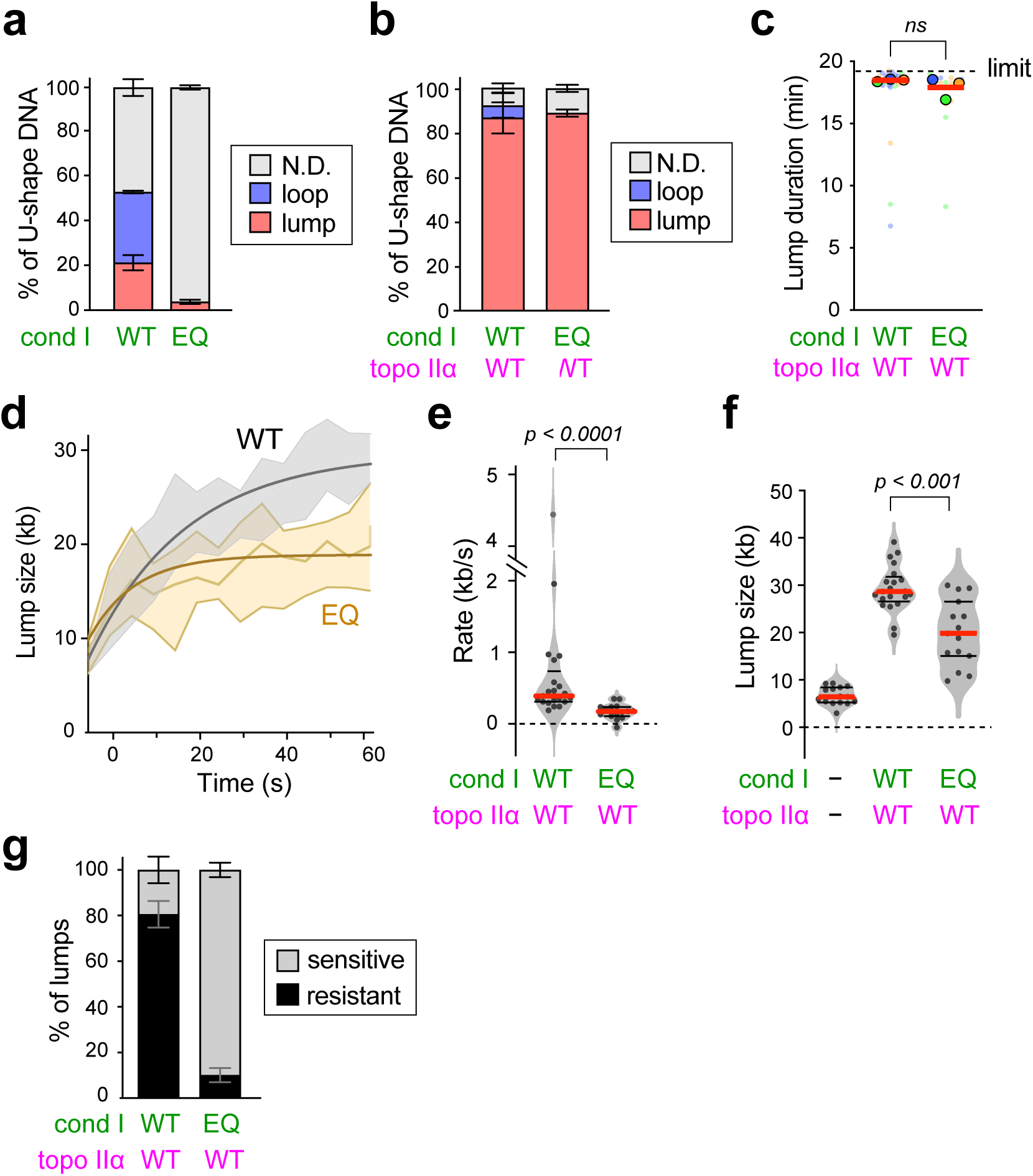
ATP hydrolysis by condensin I strongly stimulates protease-resistant lump formation. **a**, Frequency of lumps and loops on U-shape DNA formed by cond I^WT^ (WT) or cond I^EQ^ (EQ). Data were collected from 3 independent trials (cond I^WT^: 21,12, 21 DNAs; cond I^EQ^: 15, 20, 19 DNAs; mean ± SEM). **b**, Frequency of lumps and loops on U-shape DNA formed by topo IIα^WT^ in the presence of either cond I^WT^ (WT) or cond I^EQ^ (EQ). Data were collected from 3 independent trials (cond I^WT^: n = 27, 50, 30 DNAs; cond I^EQ^: n = 21, 22, 10 DNAs; mean ± SEM). **c**, Stability of lumps formed by topo IIα^WT^ in the presence of either cond I^WT^ (WT) or cond I^EQ^ (EQ). The duration of lumps was measured under imaging buffer flow. The maximum duration is the observation time 20 min. The solid point denotes the median of each condition, and the red bar represents the mean of the median. Data were collected from 3 independent trials (cond I^WT^: n = 19, 6, 6 DNAs; cond I^EQ^: n = 3, 7, 4 DNAs). The P values were assessed by a two-tailed Mann-Whitney U test. **c**, Maintaining time of lump with cond I^WT^ or cond I^EQ^ in the presence of topo IIα^WT^. The time was measured from the start of lump formation up to 20 minutes of monitoring under imaging buffer flow. The solid point denotes the median of each condition and red bar represents mean of the median. Data was collected from 3 independent trials (cond I^WT^: n = 19, 6, 6 molecules, cond I^EQ^: n = 3, 7, 4 molecules). The P values were assessed by a two-tailed Mann-Whitney U test. **d**, Time trace of the size of a lump formed by cond I^WT^ (WT) or cond I^EQ^ (EQ) in the presence of topo IIα^WT^. Exponential fitting lines with interquartile ranges are shown. Data were collected from 15 DNAs from 3 independent trials. **e**, Rate of lump formation in the presence of topo IIα^WT^ (WT) and either cond I^WT^ (WT) or cond I^EQ^ (EQ). The red and black bars denote the median and interquartile, respectively (cond I^WT^: n = 15 DNAs; cond I^EQ^: n = 15 DNAs). The P values were assessed by a two-tailed Mann-Whitney U test). **f**, Size of lumps formed in the presence or absence of either cond I^WT^ (WT) or cond I^EQ^ (EQ) and topo IIα^WT^ (WT). The red and black bars denote the median and interquartile, respectively (n = 30, 15, 15 DNAs analyzed (from left to right)). The P values were assessed by a two-tailed Mann-Whitney U test). **g**, Frequency of ProK-resistant and -sensitive lumps formed by cond I^WT^ (WT) or cond I^EQ^ (EQ) in the presence of topo IIα^WT^. Data were collected from 3 independent trials (cond I^WT^: n = 26, 45, 22 DNAs, cond I^EQ^: n = 18, 20, 9 DNAs; mean ± SEM).

Whereas condensin I^EQ^ bound to DNA as stably as condensin I^WT^ (Extended Data Fig. 7a), we noticed that condensin I^EQ^ formed lumps more slowly than condensin I^WT^ (Fig. 6d). The formation rate (Fig. 6e) and the size (Fig. 6f) of the lumps formed by condensin I^EQ^ were significantly smaller than those formed by condensin I^WT^. Furthermore, most of the lumps formed by condensin I^EQ^ in the presence topo IIα^WT^ (∼90%) were protease-sensitive (Fig. 6g). To test whether the DNA present in the residual protease-resistant lumps (∼10%) is topologically constrained, we applied a higher extension force by increasing the flow rate. We found that the residual protease-resistant lumps formed by condensin I^EQ^ and topo IIα^WT^ were fully tolerant to the high extension force (Extended Data Fig. 7b), in contrast to those formed by condensin I^WT^ and topo IIα^Y803F^ (Extended Data Fig. 6e). This observation suggests that condensin I^EQ^ can introduce DNA knots in the presence of topo IIα^WT^, but at a much lower frequency than condensin I^WT^.

## Discussion

The establishment of a chromatid reconstitution assay has allowed us to identify the minimal set of protein components required for the core reaction of mitotic chromosome assembly^7,8^. Among the six essential components identified, condensin I and topo IIα are the only two ATPases. Although the two ATPases have extensively been studied using various single-molecule assays, they have never been studied in combination. The current study represents the first attempt to study the interplay between condensin I and topo IIα at single-molecule resolution, with the aim of bridging the gap in our knowledge between the reconstitution and single-molecule assays.

Our results reported in the current study are summarized in Fig. 7a. In our current experimental setup, condensin I alone generates loops and unstable lumps on the U-shape DNA substrates (Fig. 1). We speculate that the observed unstable lumps are a precursor of the loops. Topo IIα alone, at the low concentration used in the current study, has little or no impact on the conformation of the DNA substrates. However, when condensin I and topo IIα are added together, stable lump formation dominates over loop formation (Fig. 1). Importantly, this condensin I/topo IIα-mediated lump formation depends on the CTD of topo IIα (Fig. 5), but not on its catalytic activity (Fig. 4). Although we still do not know how the condensin I/topo IIα-mediated lumps are formed, the requirement for the CTD suggests that topo IIα must stay on the DNA for a long time through CTD-mediated binding to a specific DNA structure created by condensin I (e.g., DNA crossover or bent DNA). We do not exclude the possibility that topo IIα physically interacts with condensin I on DNA, although there is no direct evidence for condensin-topo II interactions in solution (i.e., without DNA) in eukaryotic systems (e.g., Ref. 32).

**Fig. 7.**
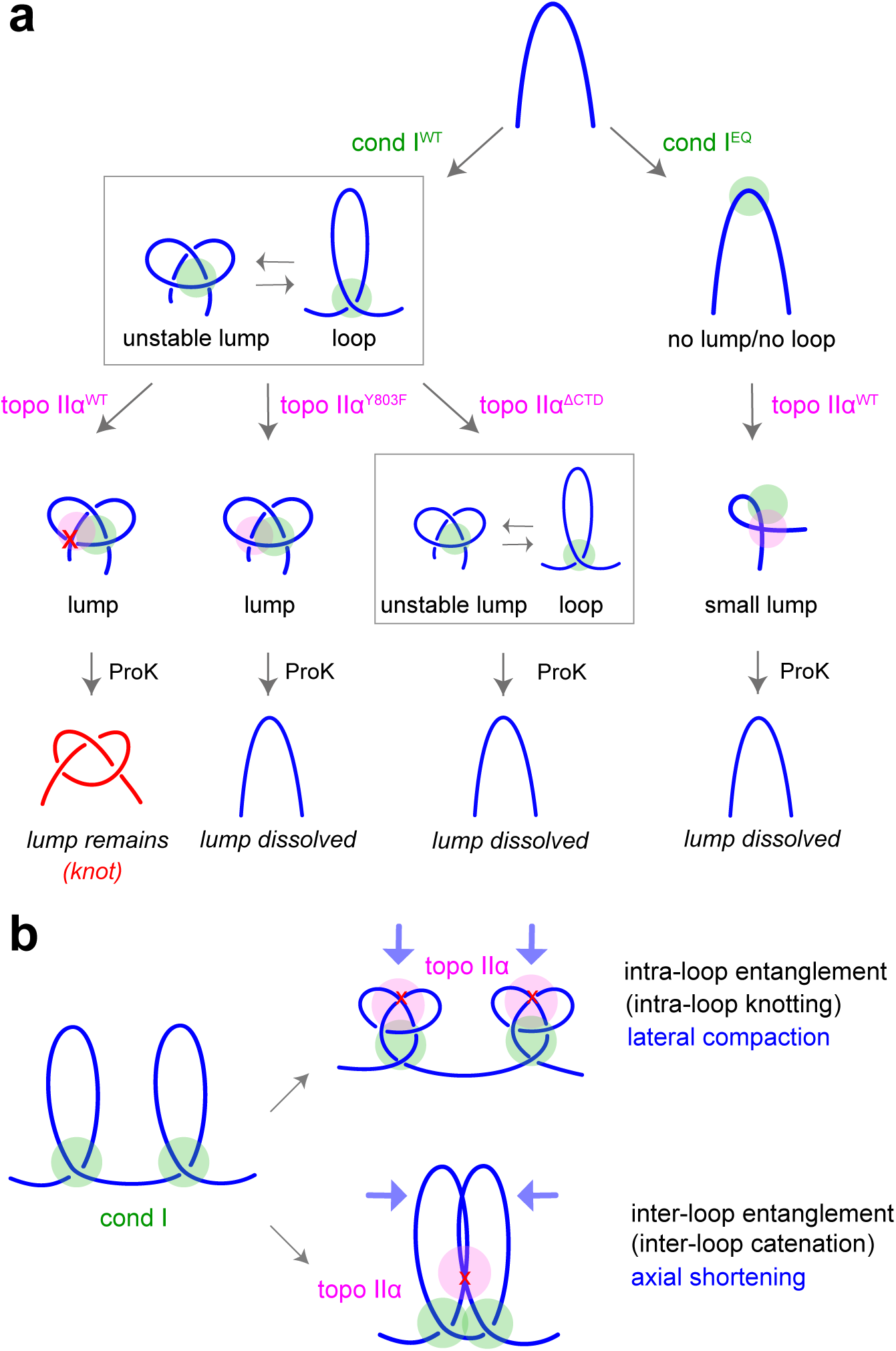
Interplay between condensin I and topo IIα. **a**, Observations in the current study. Wild-type condensin I (cond I^WT^) forms a stable lump in the presence of wild-type topo IIα (topo IIα^WT^). Stable lump formation per se does not require DNA strand passage by topo IIα (topo IIα^Y803F^), but depends on the CTD of topo IIα (topo IIα^ΔCTD^). However, the lump formed in the presence of topo IIα^WT^ becomes protease-resistant because the DNA within the lump is converted to a knotted form. A mutant condensin I complex deficient in ATP hydrolysis (cond I^EQ^), in the presence of wild-type topo IIα^WT^, forms a small lump in which DNA is rarely knotted. **b**, Functional implications in mitotic chromosome assembly. On the one hand, the entanglement within single loops (i.e., intra-loop entanglement) would facilitate lateral compaction and stabilization of mitotic chromosomes. On the other hand, the entanglement between adjacent loops (i.e., inter-loop entanglement) would promote axial shortening and stabilization of mitotic chromosomes. These two possibilities, which are not mutually exclusive, would readily occur in the highly crowded environment created during mitotic chromosome assembly.

The condensin I/topo IIα-mediated lumps become protease-resistant when topo IIα is catalytically active (Fig. 4). The following two lines of evidence strongly suggests that the protease-resistant lumps contain knotted DNA (Fig. 7a). First, the topo IIα^Y803F^ mutant supports the formation of stable lumps but fails to render them protease-resistant. Second, the protease-resistant lumps, after deproteinization, can be resolved with freshly added topo IIα. Most of the protease-resistant lumps are dissolved in a single step, suggesting that most of the knots generated under the current conditions are simple knots, such as 3-noded knots, similar to those observed in the previous bulk biochemistry using condensin I and a type II topoisomerase^13^.

A recent preprint using magnetic tweezers has reported that a low concentration (∼0.1 nM or less) of human topo IIα displays the ability to compact DNA, possibly through a mechanism of a polymer-collapse phase transition^33^. The compaction activity is fully dependent on the CTD of topo IIα, but is independent of condensin I. Thus, the topo IIα-mediated DNA compaction reported in the preprint is clearly distinct from the condensin I/topo IIα-mediated lump formation described in the current study. Nevertheless, it is important to note that both studies highlight the hitherto underappreciated role of the topo IIα CTD in the conformational changes of DNA that is separable from its strand passage activity.

How might condensin I-mediated loop extrusion be coupled to topo IIa-mediated knotting? A mutant condensin I complex harboring the ATP hydrolysis-deficient SMC subunits (condensin I^EQ^) generates lumps at the same frequency as condensin I^WT^ in the presence of topo IIα (Fig. 6). We notice that the size of condensin I^EQ^ /topo IIα-mediated lumps was substantially smaller than that of condensin I^WT^ /topo IIα-mediated lumps and that the former was far more protease-sensitive than the latter. Here we interpret our observations based on the recently updated model of condensin-mediated loop extrusion^34^. According to this model, upon ATP binding and head-head engagement, a small loop is formed in the SMC lumen (i.e., the “feeding” loop). Upon ATP hydrolysis and head-head disengagement, the feeding loop is transported into the kleisin lumen, where it is merged into the “extruding” loop (Extended Data Fig. S8). We speculate that the extruding loop generated by condensin I^WT^ would become an excellent substrate for topo IIα-mediated knotting. In contrast, condensin I^EQ^ forms a feeding loop but is unable to convert it into an extruding loop. This could explain why condensin I^EQ^ generates a small lump in which topo IIα-mediated knotting occurs only at a low frequency. The question of how the condensin-mediated loop extrusion process imposes superhelical tension on DNA has only recently been recognized in the field^29,35,36^. The currently available data are somewhat fragmentary, and our interpretation is partially, if not completely, consistent with the emerging picture. Clearly, future studies are needed to address this important question.

What are the functional implications of the current observations for mitotic chromosome assembly and stabilization? Our finding that the CTD of topo IIα is essential for the formation of protease-resistant lumps, hence DNA knotting, would provide an important clue (Fig. 1a). It has been shown that topo IIα^ΔCTD^ is proficient in topo IIα-catalyzed decatenation of circular DNA but is deficient in its efficient catenation (Fig. 1a)^8,37^. In addition, topo IIα^ΔCTD^ is proficient in inter-chromatid disentanglement but is deficient in intra-chromatid entanglement that is predicted to support chromatid thickening (axial shortening) in the reconstitution assay (Fig. 1b)^8^. Importantly, unlike topo IIα^WT^, topo IIα^ΔCTD^ is not detectable on chromosomes during the assembly processes. These observations have been interpreted to mean that the long residence time imposed by the CTD helps topo IIα to catalyze catenation^8,37^ and intra-chromatid entanglement in the reconstitution assay^8^. We speculate that the same is true for the topo IIα-stimulated lump formation and DNA knotting reported in the current study. Although direct evidence for the occurrence of intra-chromatid entanglements (self-entanglements) in mitotic chromosomes is lacking, an early study using micromanipulation approach^21^ and a recent study using high-throughput chromosome conformation capture (Hi-C) and polymer simulations^22^ have provided data consistent this idea.

We consider two scenarios for the role of intra-chromatid entanglements within mitotic chromosomes (Fig. 7b). The first is the entanglement within single loops (i.e., intra-loop entanglement), which would facilitate lateral compaction and stabilization of mitotic chromosomes. The second is the entanglement between adjacent loops (i.e., inter-loop entanglement), which would promote axial shortening and stabilization of mitotic chromosomes. The products observed in the current study (DNA knots) are roughly equivalent to those resulting from intra-loop entanglements. However, we consider that it may be possible to recapitulate inter-loop entanglements by modifying the current setup of single-molecule analyses. In any case, it is reasonable to speculate that the two reactions hypothesized here, which are not mutually exclusive, would readily occur in highly crowded environments created during mitotic chromosome assembly.

Finally, it should be added that, although the DNA present within the protease-resistant lumps is most likely a simple form of DNA knots (i.e., three-noded knots), it is uncertain whether they have specific chiralities. In this sense, it is worth mentioning that a mitotically phosphorylated form of native condensin I complex purified from *Xenopus* egg extracts generates DNA knots with a specific chirality (i.e., positive 3-noded knots) in the presence of a type II topoisomerase^13^. Because the recombinant condensin I complexes used in the current study are not subjected to mitotic phosphorylation, the knots discussed in the current study may not have specific chiralities. A better characterization of the interplay between condensins and topo IIα *in vitro* and on mitotic chromosomes in situ is an important direction in the future.

## Methods

### Preparation of recombinant condensin I holocomplexes

For the expression of recombinant condensin complexes in insect cells, the Bac-to-Bac Baculovirus Expression System (Thermo Fisher Scientific) was used as described previously^25^. The ATP hydrolysis-deficient mutations of mSMC2 (E1114Q) and mSMC4 (E1218Q) were introduced as described previously^38^. These mutations were referred to as transition-state (TR) mutations in Ref#38, whereas the same mutations are referred to as EQ mutations in the current study. Purification and labeling of Halo-tagged versions of the condensin I holocomplexes were performed as described previously^25^.

### Preparation of recombinant topo IIα proteins

Plasmid DNAs encoding *Xenopus laevis* topo IIα (topo IIα^WT^ [residues 1-1,579] and topo IIα^ΔCTD^ [residues 1-1,205]) with an N-terminal 3xFLAG-tag and a C-terminal Strep II-tag, were constructed as described previously^8^. To prepare topo IIα^Y803F^, an amino acid substitution of Y803F was introduced into the original plasmid DNAs using the QuikChange XL site-directed mutagenesis kit (Agilent, 200517). An N-terminally SNAP-tagged topo IIα construct was created with a synthetic DNA fragment encoding the SNAP tag (codon-optimized for *Trichoplusia ni*). Using these plasmid DNAs, recombinant bacmid DNAs and baculoviruses were produced as described previously^8^. Recombinant proteins were expressed in HighFive insect cells by infecting them with amplified baculovirus. For purification of the full-length versions of topo IIα (topo IIα^WT^ and topo IIα^Y803F^), cell lysate was subjected to tandem chromatography using Strep-Tactin Sepharose resin (IBA, 2-1201-010) and a Capto HiRes S 5/50 cation exchanger column (Cytiva, 29275877). For purification of topo IIα^ΔCTD^, a combination of Strep-Tactin Sepharose resin and HiTrap Q HP anion exchanger column (Cytiva, 17115301) was used. Purified SNAP-topo IIα derivatives were fluorescently labeled with the SNAP-Surface AlexaFluor 546 ligand (New England Biolabs, S9132) according to the manufacturer’s instructions. All purified topo IIα proteins were dialyzed against KHG200/50 buffer (20 mM HEPES-KOH [pH 7.7], 200 mM KCl, 50% glycerol), aliquoted, snap-frozen in liquid nitrogen, and stored at −80°C until use. The enzymatic activity of each recombinant topo IIα derivative was validated through kinetoplast DNA decatenation assays.

### Preparation of microfluidic device

Single-molecule imaging was performed in cruciform flow-cell channels. For preparation of coverslips (Matsunami 22 x 22 mm No.1-S), the coverslips were sonicated in 1 M KOH for 20 min, rinsed with water, sonicated again in ethanol, and left to stand for 1-3 nights in 2% 3-Aminopropyltriethoxysilane (Sigma Aldrich, 440140). The coverslips were then treated with 100 mM NaHCO3 containing 100 mg/ml PEG (Laysan Bio, #MPEG-SVA-5K) and BioPEG (Laysan Bio, # BIO-PEG-SVA-5K) overnight. The coverslips were then gently washed with water, dried, and stored at −80℃. Before use, the coverslips were treated with 80 μM Streptavidin (Sigma Aldrich, S4762) in ELB++ (10 mM HEPES-KOH [pH7.7], 50 mM KCl, 2.5 mM MgCl2, 1 mg/ml BSA [Sigma Aldrich, 2905-OP, OmniPur BSA, Fraction V]) for at least 30 min. Excess unbound avidin was removed by washing with water.

The cruciform flow-cell channels were developed by putting glass slides and coated coverslips with double-sided tape (Grace Bio Labs GBL620001-1EA) with a cruciform channel of 2-mm width. Inlet (Natsume KN-392-1-SP-19) and outlet tubes (Natsume KN-392-1-SP-55) were connected to each channel, and the outlet tubes were connected to a syringe pump (Harvard Apparatus Pump 11 Elite) to withdraw solutions. The channels were washed with ELB++.

Biotin-conjugation at both ends of λDNA (N3011, New England Biolabs) was performed as follows: λDNA and two biotinylated oligonucleotides, which are complementary to the *cos* sites (oligo #1, 5’-GGGCGGCGACC[biotinylated-T]-3’); oligo #2, 5’-AGGT[biotinylated-T]CGCCGCCC-3’), were phosphorylated with T4 polynucleotide kinase (M0201, New England Biolabs) in the presence of ATP at 37℃ for 3 hr. Oligo #1 was mixed with λDNA in T4 ligation buffer (B0202, New England Biolabs), annealed by cooling the mixture gradually from 65℃ to 10℃, and then ligated with T4 DNA ligase (M0202, New England Biolabs) and ATP at 16℃ for 2 hr. Finally, oligo #2 was annealed to another end of λDNA and ligated in the same way.

The resultant biotin-conjugated λDNA was injected into the perpendicular channel of the flow-cell at a final concentration of 11.5 pM in DNA buffer (50 mM Tris-HCl [pH7.5], 50 mM NaCl, 5 mM MgCl2) with either 22 nM SYTOX Orange (Thermo Fisher Scientific, S11368, in Figs. Fig. 1c-h, Fig. 2a-c, Fig. 3-6, Fig. S1-2, S3a, c, S4b, S5, S6d, e, S7) or SYTOX Green (Thermo Fisher Scientific, S7020, in Figs. 2d, g, h, S3b, and S4a), and tethered onto coverslips under the flow at 2 μl/min for 7 min. Unbound λDNAs were washed out first with DNA buffer and then with imaging buffer (IB: 50 mM Tris-HCl [pH7.5], 50 mM NaCl, 5 mM MgCl2, 3 mM ATP, 1mM DTT, 22 nM SYTOX Orange). In the experiments using Alexa546-labeled topo IIα (topo IIα^A546^), high salt-imaging buffer (50 mM Tris-HCl [pH7.5], 100 mM NaCl, 5 mM MgCl2, 3 mM ATP, 1 mM DTT, 22 nM SYTOX Green) was used to prevent non-specific binding of topo IIα^A546^(Fig. 2d, g, h, Fig. S1d, Fig. S4a). To observe U-shape DNA, the flow direction was switched from the perpendicular to the horizontal channel at a flow rate of 20 μl/min (estimated tension is < 0.2 pN).

### Lump formation assay

Condensin I and unlabeled topo IIα were simultaneously introduced into U-shape DNA-tethered flow cells with imaging buffer at a flow rate of 20 μl/min for 100 sec. Condensin I was introduced at 1 nM in all experiments and topo IIα was introduced at either 125 pM (Fig. 1c-h, Fig. 2 a-c except sample without ATP, Fig. 36, Fig. S1, Fig. S2, Fig. S3a, c, S6d, S7a) or 88 pM (Fig. 2 a-c, sample without ATP, Extended Data Fig. 4b, Fig. 5, Fig. 6e, Fig.S7b). Unbound proteins were washed out with imaging buffer. When ATP dependency was tested, we used imaging buffer without ATP instead (Fig. 2a-c, Fig. 3c). When both condensin I^A488^ and topo IIα^A546^ were visualized, due to the overlap of the fluorescent wavelengths of Alexa488 and SYTOX Green, DNA was first detected by SYTOX Green to identify the places of U-shape DNA. After washing out SYTOX dye, DNA binding of condensin I ^A488^ and topo IIα^A546^ was monitored in real time. At the end of monitoring, DNA was stained again with SYTOX Green to confirm lump formation. To test protease resistance, lumps were treated with 0.5 mg/ml Proteinase K (ProK, Roche, #03115873001) in imaging buffer at a flow rate of 20 μl/min. When topo IIα^A546^ was visualized, the lumps were treated by ProK after the 2nd SYTOX staining (Fig. S4a). In topo IIα reinjection assay (Fig. 4g), ProK was washed out with imaging buffer at 20 μl/min for 100 sec, and then 250 pM topo IIα was reinjected with imaging buffer at 20 μl/min. In the intense wash assay (Extended Data Figs. 6e and 7b), after washing out ProK, the flow rate was increased from 20 μl/min to 1 ml/min for 50 sec, and then decreased to 20 μl/min again to see whether or not the lumps remained. To calculate the number of folds of DNA intensity, protease-resistant lumps were formed as described above, then the flow rate was increased in a stepwise manner: at 20, 100, 200, 400 μl/min for 100 sec, then at 1 ml/min for 25 sec, and finally at 20 μl/min for 100 sec.

In all imaging experiments except the bleaching assay, condensin I^A488^ and SYTOX Green-stained DNA were visualized using a 350 μW 488-nm laser (Coherent Sapphire LP 488±2, 150 mW) and topo IIα^A546^ and SYTOX Orange-stained DNA were visualized using a 100 μW 561-nm laser (Coherent Sapphire LPX 561±2, 480 mW). All assays were performed at room temperature.

### Bleaching assay

For the bleaching assay to detect single-molecule intensities of condensin I^A488^ and topo IIα^A546^ (Figs. 2e-f), 1 nM condensin I^A488^ or 75 pM topo IIα^A546^ was non-specifically attached to coverslips. The fluorescently labeled proteins were then bleached by either a 3.5 mW 488-nm laser or 5.0 mW 561-nm laser, the images were taken every 0.5 sec, and the bleaching step was detected. The step size was determined by calculating the difference in average intensity of five flames before and after the bleaching.

### DNA decatenation assay

One hundred nanograms of a catenated DNA substrate (kinetoplast DNA; TG2013, TopoGEN) were mixed with 40 ng of recombinant topo IIα in a 10-µl volume (20 mM HEPES-KOH [pH 7.7], 80 mM KCl, 5 mM MgCl2, and 2 mM ATP), and incubated at 22°C. At various times, aliquots were taken and treated with SDS (0.5%) and proteinase K (1.0 mg/ml; P-4032, Sigma) at 37°C for 1 h. The resultant DNAs were purified with phenol and separated by gel electrophoresis on a 0.8% agarose gel in TAE. After being stained with ethidium bromide, fluorescent images were acquired with an image analyzer (Amersham Imager 680 [version 2.0.0], Cytiva).

### Depletion and add-back assays using recombinant topo IIα in *Xenopus* egg extracts

The high-speed supernatant of metaphase-arrested *Xenopus* egg extracts (M-HSS) and demembranated *Xenopus* sperm nuclei were prepared as described previously^39,40^. A rabbit polyclonal antibody was raised against a synthetic peptide corresponding to the C-terminal amino acid sequence of *Xenopus laevis* topo IIα (GRQKKPVTYLEDSDDDF) and affinity-purified (referred to here as anti-topo IIα, in-house identifier AfR474-2). One hundred microliters of M-HSS were incubated with Dynabeads Protein A (Thermo Fisher Scientific, 10002D) conjugated with 20 µg of anti-topo IIα. After two successive rounds of incubation on ice (30 min each), the supernatants were isolated from the beads under a magnetic field and used as topo IIα-depleted M-HSS for subsequent assays. Control mock-depleted M-HSS was prepared using non-immune rabbit IgG (Sigma-Aldrich, I-5006) instead of anti-topo IIα. To assess the ability of recombinant topo IIα (wild-type and Y803F mutant) to support mitotic chromatid assembly in egg extracts, sperm nuclei were incubated with either mock-depleted or topo IIα-depleted M-HSS that had been supplemented with buffer (KHG200/50) or recombinant topo IIα at 22°C for 120 min. The resultant chromatin structures were fixed, stained with DAPI, labeled with anti-topo II monoclonal antibody (M052-3, MBL, RRID: AB_592894), and analyzed by fluorescence microscopy as described previously^8^

### Immunofluorescence labeling of FLAG-topo IIα

After lump formation by condensin I and FLAG-tagged topo IIα, the lumps were washed with imaging buffer in the presence or absence of ProK for 100 sec, and then ProK was washed out by another 100-sec wash. The channels were washed with ELB++ at 20 μl/min for 5 min, 1 μg/ml anti-FLAG M2 antibodies (Sigma Aldrich, F1804) in ELB++ was introduced at 10 μl/min for 18 min, and washed with ELB++ at 20 μl/min for 5 min. Then 2 μg/ml Alexa Fluor 647-conjugated anti-mouse IgG antibodies (Thermo Fisher Scientific, A-21235) in ELB++ were introduced at 10 μl/min for 10 min and washed with ELB++ at 20 μl/min for 5 min. Finally, DNAs were visualized with 11 nM SYTOX Green in ELB++, and the localization and intensity of FLAG-topo IIα on the lump were analyzed.

### Data collection and processing

Experiments were performed on a total internal reflection fluorescence microscope (Nikon Ti) and images were acquired every 0.5 or 5 sec using a CCD camera (Andor DU-888 X-7952) under the control of NIS software. All images were analyzed by Fiji. The intensities of condensin I and topo IIα on all U-shape DNA were measured, and only those with intensities equivalent to a single molecule were used for the analyses of lump formation. Lumps and loops on U-shape DNA were classified by visual observation. Lump stability was defined as a time from the initiation of lump formation to either the disappearance of lumps or the end of the observation (up to 20 min) at 20 μl/min. The fluorescent intensities of DNA-bound condensin I and topo IIα were measured every 5 sec and the background was subtracted. The lump size was calculated by normalizing the intensity of lumps to the intensity of full-length λDNA and converting it to the DNA length by multiplying 48.5 kbp. The tip of U-shape DNA, where a lump is typically formed, exhibits naturally higher DNA intensity compared to that of single dsDNA on the side arm of U-shape DNA. Hence, the size of tip without protein was also calculated for comparison. The lump rate and loop rate were determined based on the average formation rate as long as they were glowing. Relative DNA length was calculated by dividing the DNA length in each frame by the average DNA length from the first five frames taken before protein injection. To determine the number of folds of lump intensities, the line profiles were taken from cross-section DNA intensities on tip DNA with or without lump or on side DNA. In the resulting line profiles, the relative sum intensities (Area Under the Curve, AUC) and the relative maximum DNA intensities (Max) were calculated by subtracting the background from the integral of tip and side arm DNA profiles. All images shown in the figures were background subtracted, smoothed, and resized for clarity.

## Acknowledgments

We are grateful to Keiko Kimura and Mai Shimoura for the preparation of the flow-cell devices, Ryota Sakata and Yoshimi Kinoshita for technical advice on microscopic work, and Makoto M. Yoshida for discussions. The work in the authors’ laboratory was supported by JSPS Grants-in-Aid for Scientific Research/KAKENHI (#19H05755 and #22H02551 to K.S.; #23K05649 to K.K.; #18H05276 and #20H05938 to T.H.; #20H05933 and #20H05937 to T.N.). Y.T was supported by Grant-in-Aid for JSPS Fellows.

## Extended Data Figure legends

**Extended Data Fig. 1.**
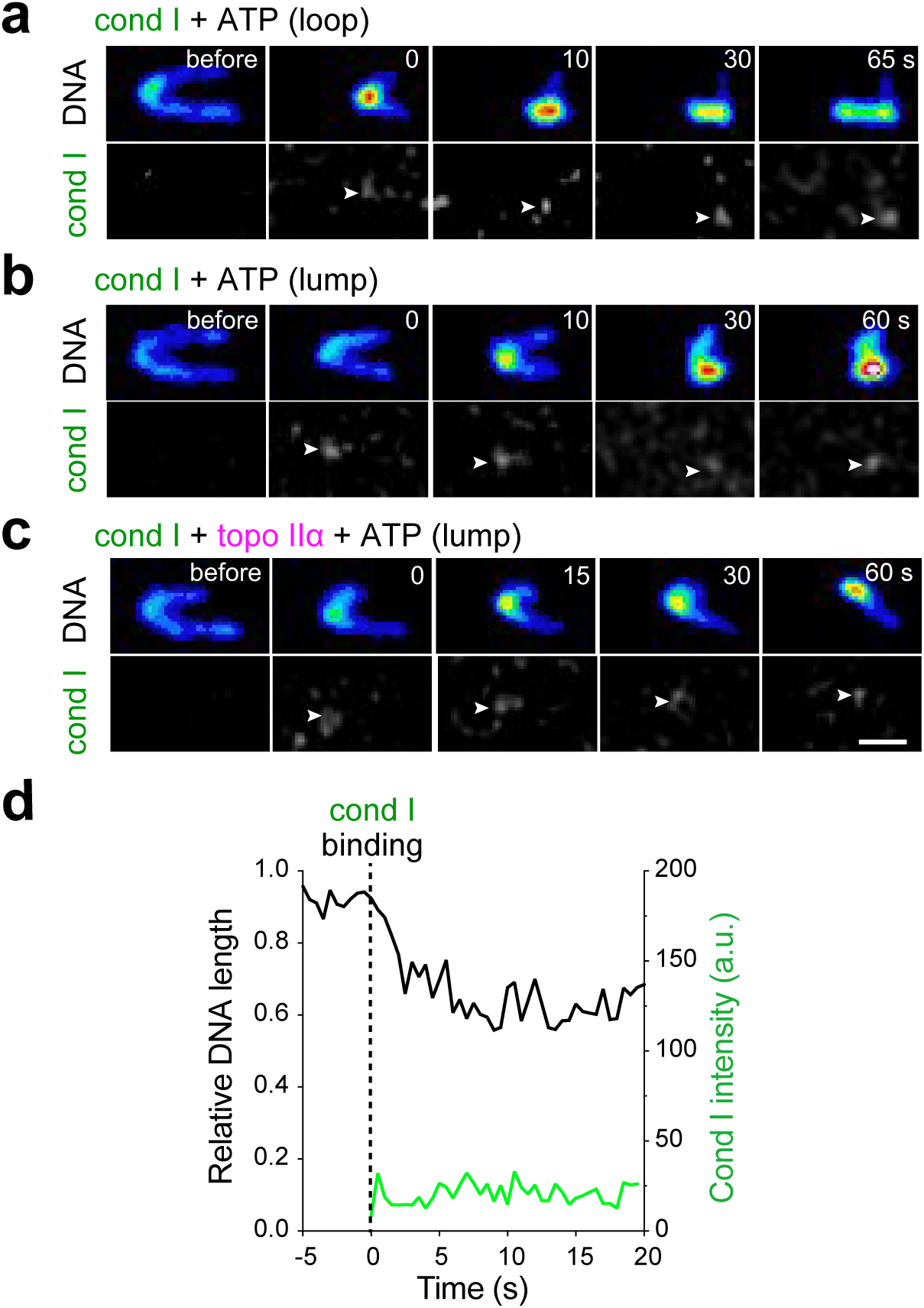
Time-lapse imaging of loop and lump formation. **a-c**, Snapshots of loop or lump formation with single condensin I (cond I) complexes in the presence or absence of topo IIα. All reactions contained ATP. Time after cond I binding is indicated. Bar: 2 μm. **d**, Time trace of DNA length and condensin I (cond I) intensity in the presence of cond I and topo IIα (with ATP). DNA length was shown as a relative DNA length to that before cond I and topo IIα addition.

**Extended Data Fig. 2.**
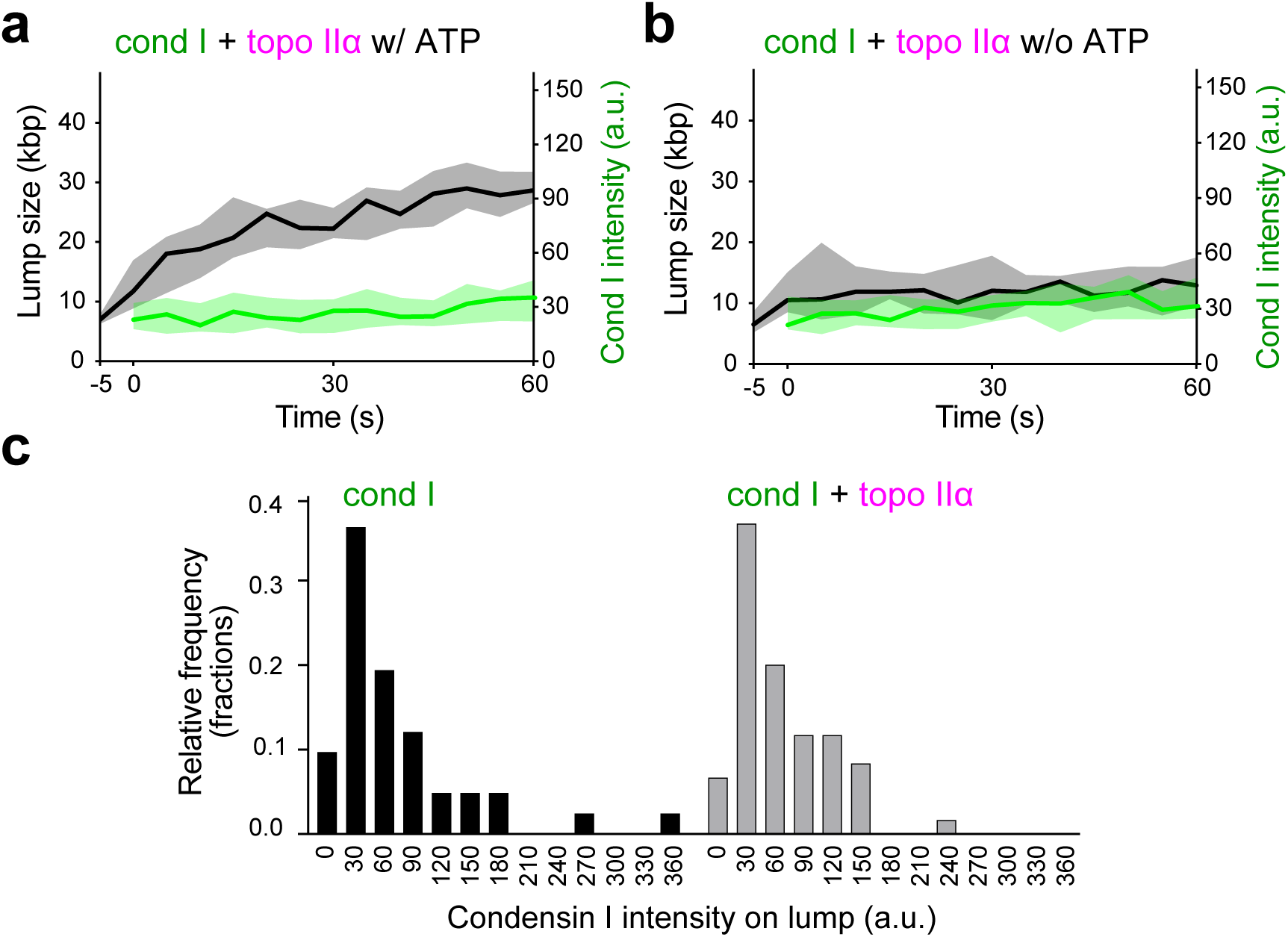
Single condensin I complexes form DNA lumps in an ATP-dependent manner. **a-b**, Time trace of the cond I^Alexa488^ intensity and the lump size formed by cond I and topo IIα in the presence (a) or absence (b) of ATP. Time 0 indicates the time when cond I first bound to the DNA. Data were collected from 20 samples from 2 independent trials (a) or 15 samples from 3 independent trials (d). Median and interquartile range are shown. **c**, Fluorescence intensity distribution of cond I^A488^ on lumps in the presence or absence of topo IIα. Data were collected from more than 40 lumps per condition.

**Extended Data Fig. 3.**
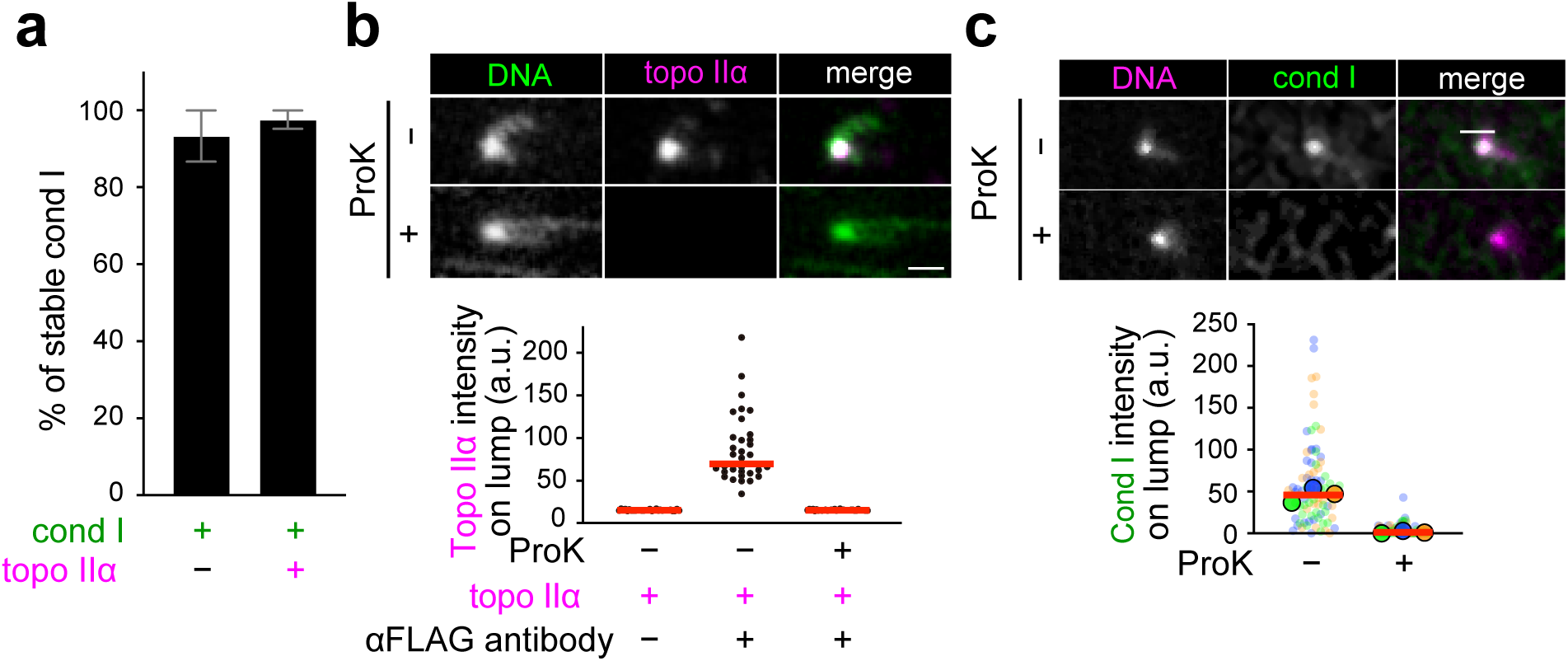
Characterization of condensin I/topo IIα-dependent lump. **a,** Percentage of condensin I (cond I) stably bound to DNA throughout 20-min observations. Data were collected from 3 independent trials (cond I alone: n = 5, 4, 6 DNAs; cond I and topo IIα: n = 13, 14, 16 DNAs; mean ± SEM). **b**, Immunofluorescence labeling of FLAG-topo IIα on a lump with or without proteinase K (ProK) treatment. FLAG-topo IIα was labeled with an anti-FLAG antibody and DNA was counterstained with SYTOX Green (upper panel). The signal intensities were quantified (lower panel). The red bars represent the median. Data were collected from 1 trial. n = 30, 36, 34 (from left to right). **c**, Fluorescence intensity of cond I^A488^ on a lump before (-) or after (+) ProK treatment. DNA was counterstained with SYTOX Orange (upper panel). The signal intensities were quantified (lower panel). The solid points represent the median of each condition, and the red bars represent the mean of the median. Data were collected from 3 independent trials (ProK (-): n = 29, 28, 29 DNAs; ProK (+): n = 29, 28, 29 DNAs).

**Extended Data Fig. 4.**
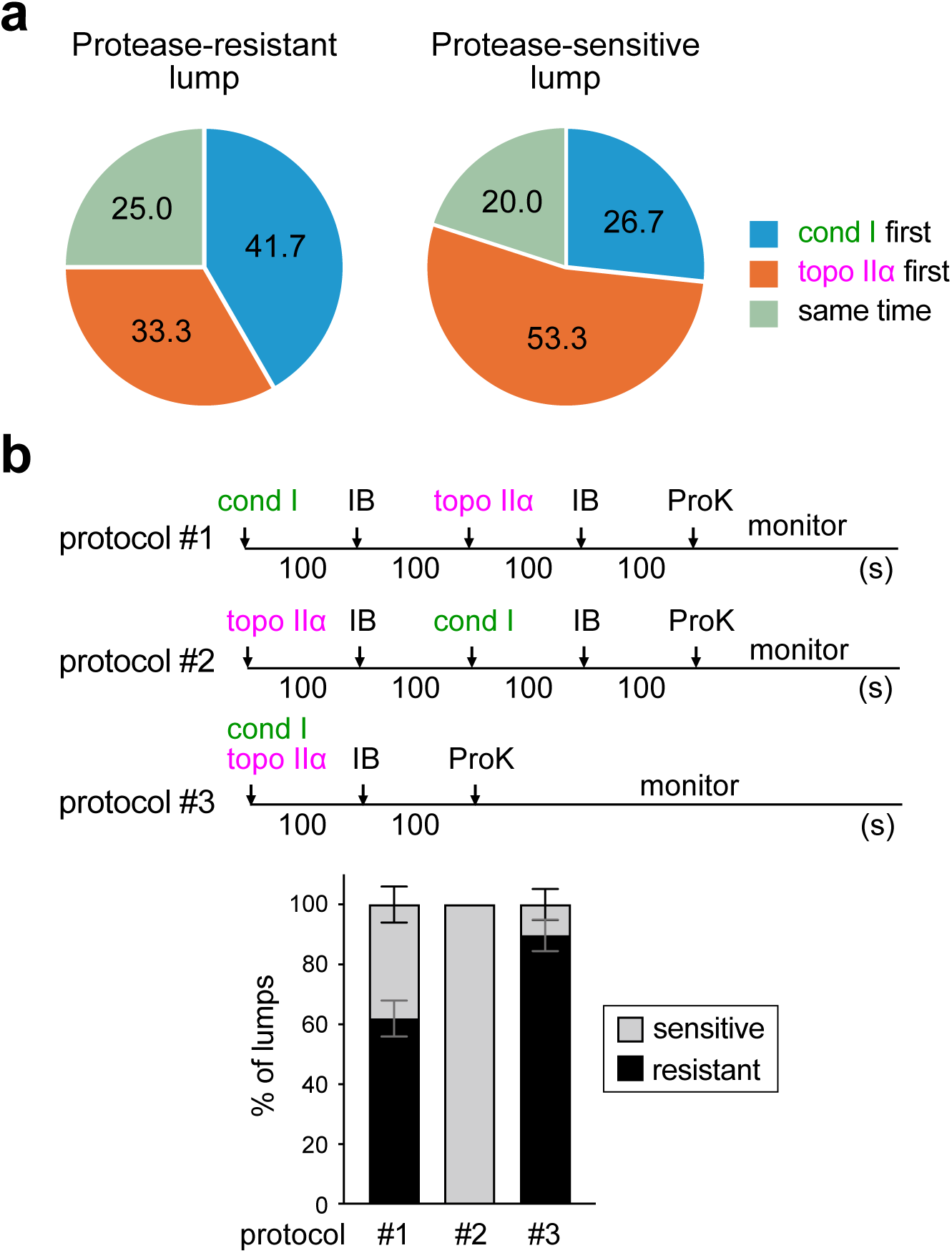
Kinetics of condensin I and topo IIα binding to DNA. **a**, Individual cond I^A488^ and topo IIα^A546^ signals were tracked during lump formation and categorized based on their orders to bind to DNA. Of the 15 protease-sensitive cases, 4 had cond I binding first (blue), 8 had topo IIα binding first (orange), 3 had both cond I and topo IIα binding simultaneously (right green). Of the 12 protease-resistant cases, 5 had cond I binding first (blue), 4 had topo IIα binding first (orange), 3 had both cond I and topo IIα binding simultaneously (right green). **b**, Frequency of protease-resistant lump formation in the experiment using 3 different protocols (mean ± SEM). In protocol #1, cond I was first injected for 100 sec, washed out for 100 sec, and then topo IIα was injected for 100 sec. In protocol #2, topo IIα was injected for 100 sec, washed out for 100 sec, and then cond I was injected for 100 sec. In protocol #3, cond I and topo IIα were injected simultaneously. Data were collected from 3 independent trials (protocol #1: n = 11, 5, 15 DNAs; protocol #2: n = 14, 9, 23 DNAs; protocol #3: n = 7, 5, 8 DNAs). The P values were assessed by a two-tailed Mann-Whitney U test.

**Extended Data Fig. 5.**
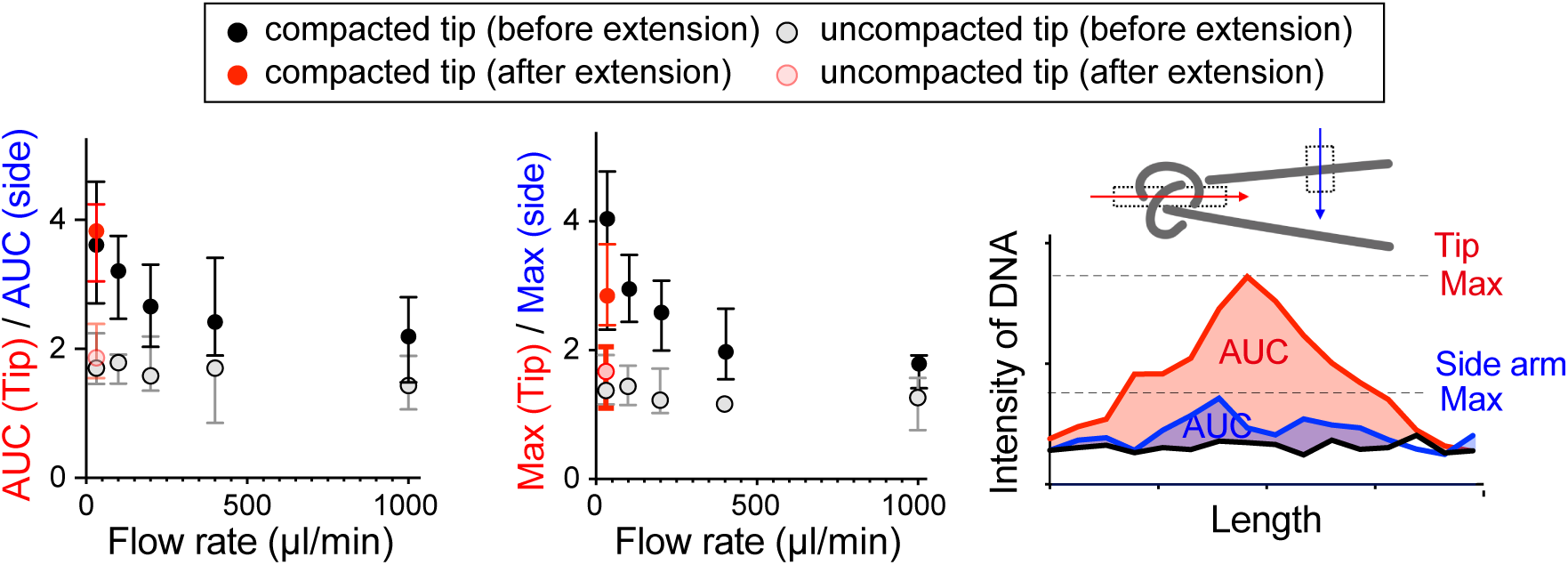
Protease-resistant lumps are resistant to DNA extension. Protease-resistant lumps were formed in the presence of condensin I and topo IIα, DNA was stained with SYTOX Orange, and the cross section of DNA was traced either on the tips of U-shape DNA or side arms (as controls) under different flow rates (20, 100, 200, 400, and 1000 μl/min). From the resulting line profiles (an example is shown on right) of the tip and the side arm DNA, the relative sum intensities (Area Under the Curve, AUC) and the relative maximum DNA intensities (Max) were calculated by subtracting the background from the integral of tip and side arm DNA profiles.

**Extended Data Fig. 6.**
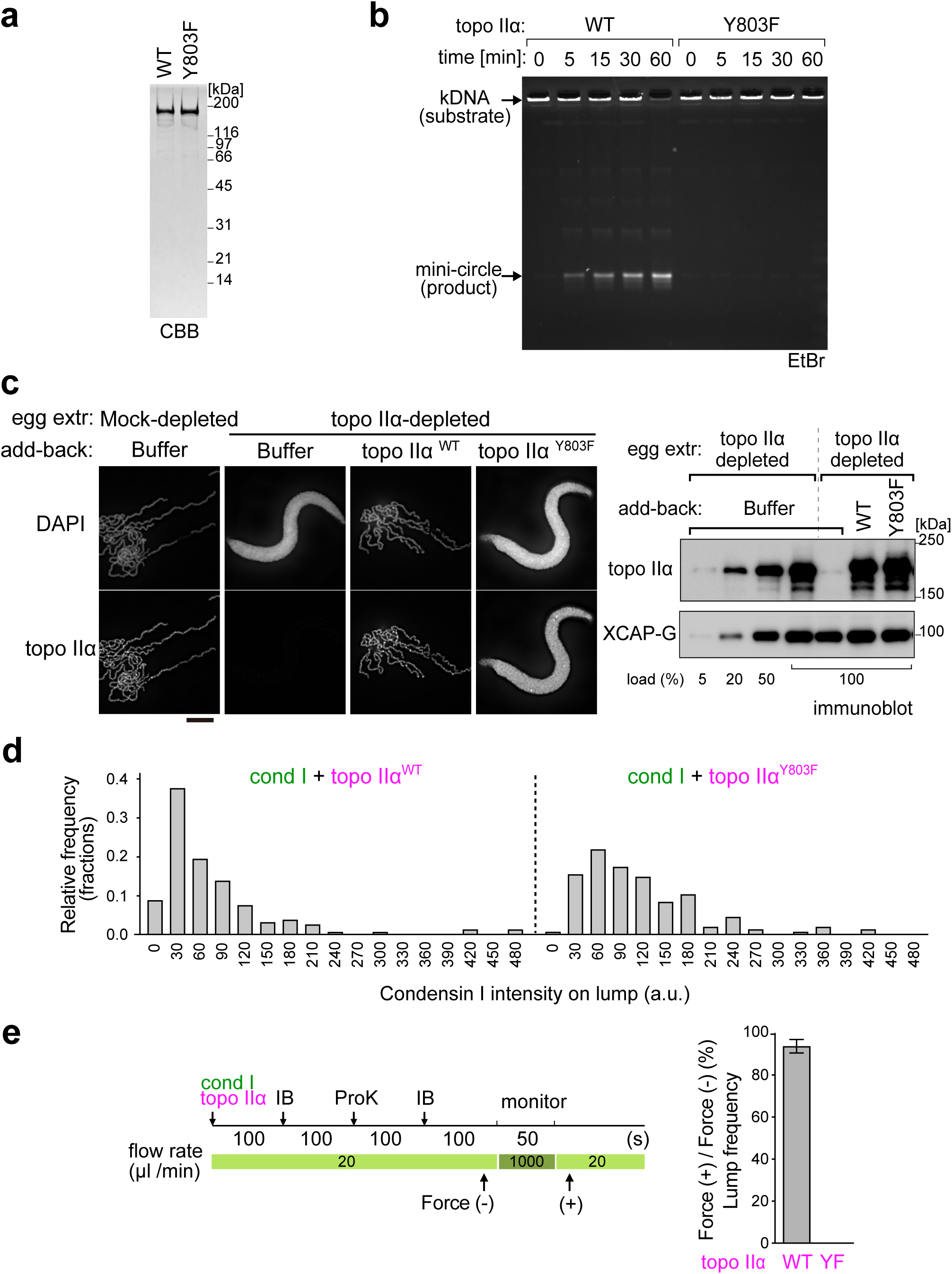
Purification and functional characterization of topo IIα^Y803F^. **a**, Recombinant *Xenopus* topo IIα wild-type (WT) and Y803F were analyzed by SDS-PAGE and stained with Coomassie Brilliant Blue (CBB). **b**, Kinetoplast catenated DNA (kDNA) was incubated with topo IIα^WT^ or topo IIα^Y803F^ in the presence of ATP at 22°C. The resultant DNAs were recovered at the indicated time points, purified, and analyzed by agarose gel electrophoresis. The gel was stained with ethidium bromide. **c**, *Xenopus* sperm nuclei were incubated in a control extract (mock-depleted) or an extract depleted of topo IIα that had been supplemented with buffer, topo IIα^WT^, or topo IIα^Y803F^. After a 120-min incubation at 22°C, the resultant structures were fixed and labeled with anti-topo IIα antibody. DNA was counterstained with DAPI. Bar, 5 µm (left). To estimate the efficiency of depletion and add-back, an aliquot of each extract, alongside decreasing amounts of the mock-depleted extract, was analyzed by immunoblotting with the anti-topo IIα antibody. As a loading control, XCAP-G, a condensin I subunit, was also examined. **d**, Fluorescence intensity distribution of condensin I (cond I)^A488^ on the lumps formed by topo IIα^WT^ or topo IIα^Y803F^. Data were collected from more than 43 lumps per condition. **e**, Percentages of protease-resistant lump remaining after DNA extension. Protease-resistant lumps were formed by condensin I (cond I) and either topo IIα^WT^ (WT) or topo IIα^Y803F^ (YF). The lumps were then extended by increasing the flow rate from 20 μl/min to 1000 μl/min. Percentages of protease-resistant lumps were shown before the increase (Force (-)) and after the decrease (Force (+)) of flow rate. Data were collected from 3 independent trials (topo IIα^WT^: n = 19, 24, 5; topo IIα^Y803F^: n = 1, 1, 1, mean ± SEM).

**Extended Data Fig. 7.**
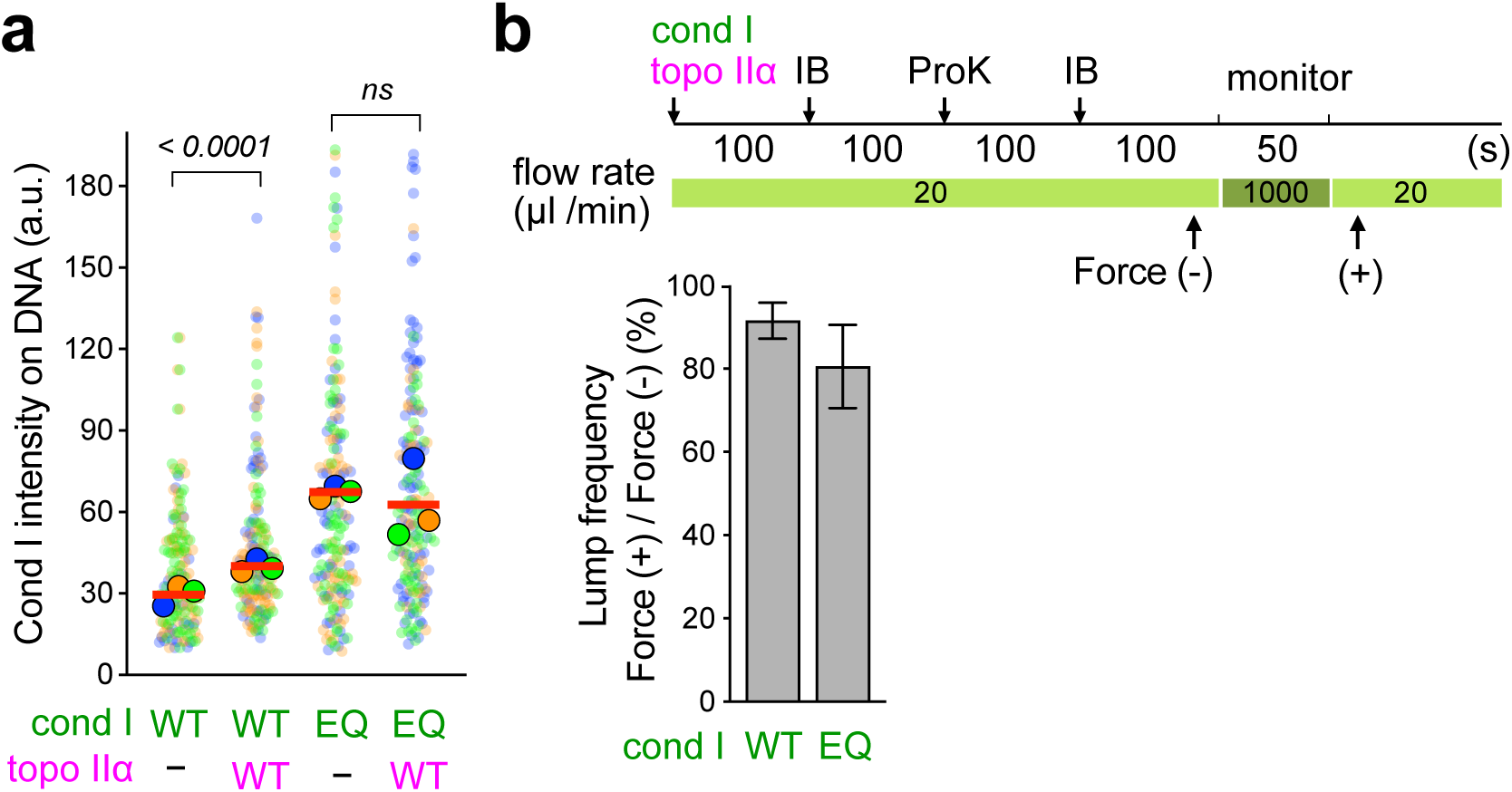
Condensin I^EQ^ has the ability to form protease-resistant lumps. **a**, Intensity of cond I^WT^ and cond I^EQ^ on U-shape DNA. The solid points represent the median of each condition, and the red bars represent the mean of the median. Data were collected from 3 independent trials (cond I^WT^ alone: n = 28, 68, 69 DNAs; cond I^WT^ and topo IIα: n = 52, 73, 48 DNAs; cond I^EQ^ alone: n = 57, 58, 61 DNAs; cond I^EQ^ and topo IIα: n = 38, 92, 63 DNAs). The P values were assessed by a two-tailed Mann-Whitney U test. **b**, Percentages of protease-resistant lumps remaining after DNA extension (mean ± SEM). Protease-resistant lumps were formed by topo IIα and either cond I^WT^ or cond I^EQ^. Lumped DNAs were then extended by increasing the flow rate from 20 to 1000 μl/min. Percentages of protease-resistant lumps were measured before the increase (Force (-)) and after the decrease (Force (+)) of the flow rate. Data were collected from 3 independent trials (cond I^WT^: n = 7, 9, 7 DNAs; cond I^EQ^: n = 3, 4, 3 DNAs).

**Extended Data Fig. 8.**
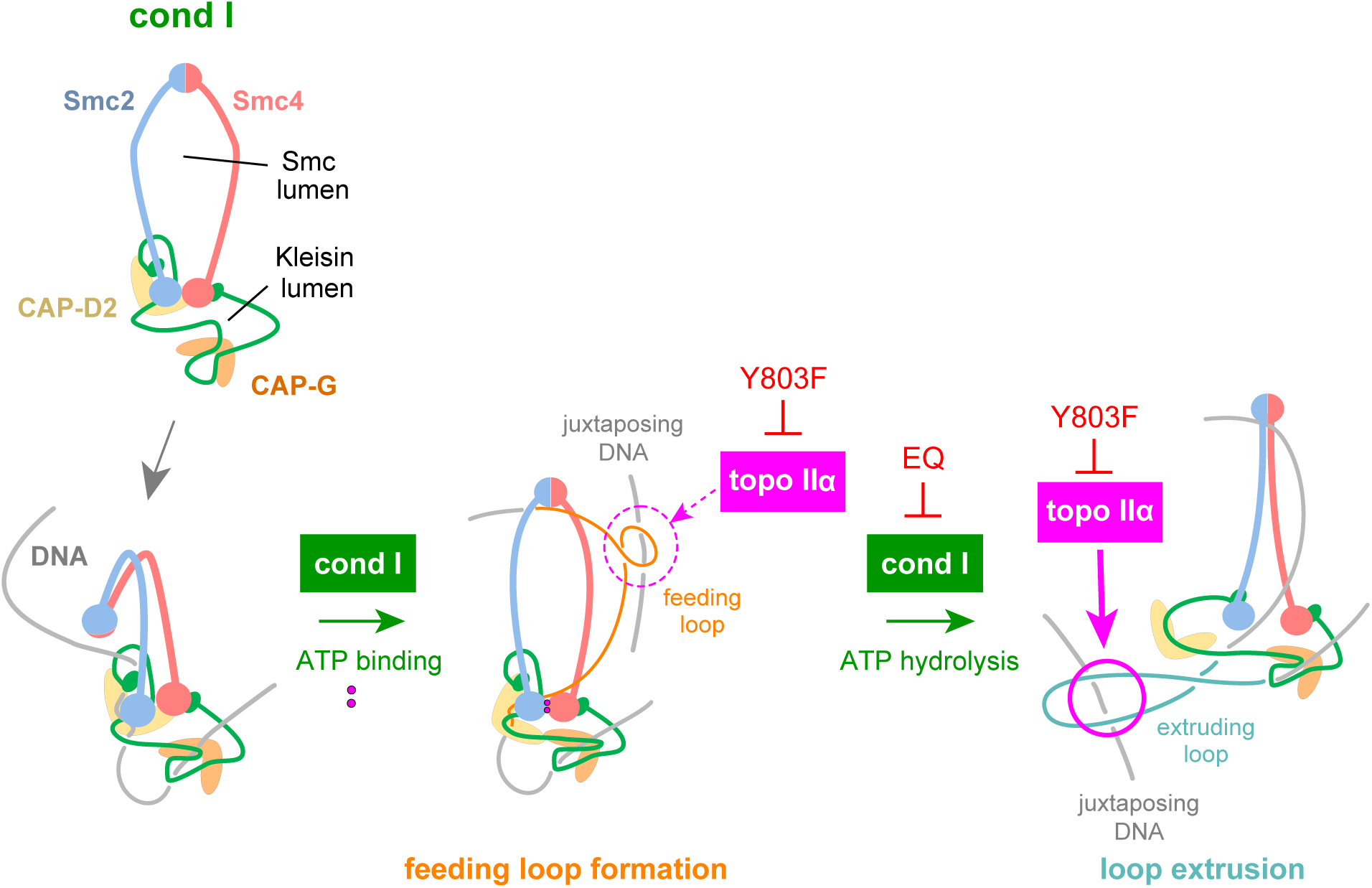
A possible model of how condensin I and topo IIα cooperate to introduce a DNA knot. DNA loops can be formed by condensin I (cond I) at the SMC lumen and at the kleisin lumen. The magenta circles indicate where the looped DNA can be crossed with its neighboring DNA. The Y803F mutation impairs the catalytic activity of topo IIα, whereas the EQ mutation impairs ATP hydrolysis by cond I. It is hypothesized here that the “extruding” loop is the major target of strand passage that introduces a DNA knot. The “feeding” loop may also be a target of strand passage at a lower frequency.

## Supplemental Movie legends

**Supplemental movie 1** An example of lump formation on DNA in the presence of condensin I (cond I) and topo IIα. The lump was formed by a single condensin I complex, and it remained stable even after ProK treatment. The arrows denote bound condensin I. Images were acquired every 5 sec. Bar: 2 μm.

**Supplemental movie 2** An example of lump formation from a DNA loop in the presence of condensin I (cond I) and topo IIα. This event occurred only when additional condensin I was bound to the pre-formed loop. The arrows denote DNA-bound condensin I. Images were acquired every 5 sec. Bar: 2 μm.

**Supplemental movie 3** An example of lump formation on DNA in the presence of condensin I alone. The lump was formed by a single condensin I complex and it was immediately resolved after ProK injection. The arrows denote DNA-bound condensin I. Images were acquired every 5 sec. Bar: 2 μm.

## References

1. Hirano, T. Condensin-Based Chromosome Organization from Bacteria to Vertebrates. Cell 164, 847–857 (2016).

2. Paulson, J.R., Hudson, D.F., Cisneros-Soberanis, F. & Earnshaw, W.C. Mitotic chromosomes. Seminars in Cell & Developmental Biology 117, 7–29 (2021).

3. Earnshaw, W.C., Halligan, B., Cooke, C.A., Heck, M.M. & Liu, L.F. Topoisomerase II is a structural component of mitotic chromosome scaffolds. J Cell Biol 100, 1706–1715 (1985).

4. Gasser, S.M., Laroche, T., Falquet, J., Boy de la Tour, E. & Laemmli, U.K. Metaphase chromosome structure. Involvement of topoisomerase II. J Mol Biol 188, 613–629 (1986).

5. Saitoh, N., Goldberg, I.G., Wood, E.R. & Earnshaw, W.C. ScII: an abundant chromosome scaffold protein is a member of a family of putative ATPases with an unusual predicted tertiary structure. J Cell Biol 127, 303–318 (1994).

6. Maeshima, K. & Laemmli, U.K. A two-step scaffolding model for mitotic chromosome assembly. Dev Cell 4, 467–480 (2003).

7. Shintomi, K., Takahashi, T.S. & Hirano, T. Reconstitution of mitotic chromatids with a minimum set of purified factors. Nat Cell Biol 17, 1014–1023 (2015).

8. Shintomi, K. & Hirano, T. Guiding functions of the C-terminal domain of topoisomerase IIalpha advance mitotic chromosome assembly. Nat Commun 12, 2917 (2021).

9. Nitiss, J.L. DNA topoisomerase II and its growing repertoire of biological functions. Nat Rev Cancer 9, 327–337 (2009).

10. Vos, S.M., Tretter, E.M., Schmidt, B.H. & Berger, J.M. All tangled up: how cells direct, manage and exploit topoisomerase function. Nat Rev Mol Cell Biol 12, 827–841 (2011).

11. Pommier, Y., Nussenzweig, A., Takeda, S. & Caroline, A. Human topoisomerases and their roles in genome stability and organization. Nature Reviews Molecular Cell Biology 23, 407–427 (2022).

12. Kimura, K. & Hirano, T. ATP-dependent positive supercoiling of DNA by 13S condensin: a biochemical implication for chromosome condensation. Cell 90, 625–634 (1997).

13. Kimura, K., Valentin, V.R., Crisona, N.J., Hirano, T. & Cozzarelli, N.R. 13S Condensin Actively Reconfigures DNA by Introducing Global Positive Writhe: Implications for Chromosome Condensation. Cell 98, 239–248 (1999).

14. Kimura, K., Hirano, M., Kobayashi, R. & Hirano, T. Phosphorylation and activation of 13S condensin by Cdc2 in vitro. Science 282, 487–490 (1998).

15. Ganji, M. et al. Real-time imaging of DNA loop extrusion by condensin. Science 360, 102–105 (2018).

16. Kong, M. et al. Human Condensin I and II Drive Extensive ATP-Dependent Compaction of Nucleosome-Bound DNA. Mol Cell 79, 99–114 (2020).

17. Goloborodko, A., Imakaev, M.V., Marko, J.F. & Mirny, L. Compaction and segregation of sister chromatids via active loop extrusion. Elife 5, e14864 (2016).

18. Postow, L., Crisona, N.J., Peter, B.J., Hardy, C.D. & Cozzarelli, N.R. Topological challenges to DNA replication: conformations at the fork. Proc Natl Acad Sci U S A 98, 8219–8226 (2001).

19. Baxter, J. et al. Positive supercoiling of mitotic DNA drives decatenation by topoisomerase II in eukaryotes. Science 331, 1328–1332 (2011).

20. Dyson, S., Segura, J., Martinez-Garcia, B., Valdes, A. & Roca, J. Condensin minimizes topoisomerase II-mediated entanglements of DNA in vivo. EMBO J 40, e105393 (2021).

21. Kawamura, R. et al. Mitotic chromosomes are constrained by topoisomerase II-sensitive DNA entanglements. J Cell Biol 188, 653–663 (2010).

22. Hildebrand, E.M. et al. Mitotic chromosomes are self-entangled and disentangle through a topoisomerase-II-dependent two-stage exit from mitosis. Mol Cell 84, 1422–1441 (2024).

23. Baxter, J. & Aragon, L. A model for chromosome condensation based on the interplay between condensin and topoisomerase II. Trends Genet 28, 110–117 (2012).

24. Sakata, R. et al. Opening of cohesin’s SMC ring is essential for timely DNA replication and DNA loop formation. Cell Rep 35, 108999 (2021).

25. Kinoshita, K. et al. A loop extrusion-independent mechanism contributes to condensin I-mediated chromosome shaping. J Cell Biol 221, e202109016 (2022).

26. Tane, S. et al. Cell cycle-specific loading of condensin I is regulated by the N-terminal tail of its kleisin subunit. Elife 11, e84694 (2022).

27. Berger, J.M., Gamblin, S.J., Harrison, S.C. & Wang, J.C. Structure and mechanism of DNA topoisomerase II. Nature 379, 225–232 (1996).

28. Liu, Q. & Wang, J.C. Identification of active site residues in the “GyrA” half of yeast DNA topoisomerase II. J Biol Chem 273, 20252–20260 (1998).

29. Martinez-Garcia, B. et al. Condensin pinches a short negatively supercoiled DNA loop during each round of ATP usage. EMBO J 42, e111913 (2023).

30. Davidson, I.F. et al. DNA loop extrusion by human cohesin. Science 366, 1338–1345 (2019).

31. Pradhan, B. et al. The Smc5/6 complex is a DNA loop-extruding motor. Nature 616, 843–848 (2023).

32. Ryu, J.K. et al. The condensin holocomplex cycles dynamically between open and collapsed states. Nat Struct Mol Biol 27, 1134–1141 (2020).

33. Wu, M., et al. Human Topoisomerase IIalpha Promotes Chromatin Condensation Via a Phase Transition. *bioRxiv*, DOI: 10.1101/2024.10.15.618281 (2024).

34. Dekker, C., Christian, H.H., Peters, J.-M. & Rowland, B.D. How do molecular motors fold the genome? Science 382, 646–648 (2023).

35. Davidson, I.F., et al. Cohesin supercoils DNA during loop extrusion. *bioRxiv*, DOI: 10.1101/2024.03.22.586228. (2024).

36. Janissen, R., et al. All eukaryotic SMC proteins induce a twist of −0.6 at each DNA-loop-extrusion step. *bioRxiv*, DOI: 10.1101/2024.03.22.586328 (2024).

37. Kawano, S. et al. DNA-binding activity of rat DNA topoisomerase II alpha C-terminal domain contributes to efficient DNA catenation in vitro. J Biochem 159, 363–369 (2016).

38. Kinoshita, K., Kobayashi, T.J. & Hirano, T. Balancing acts of two HEAT subunits of condensin I support dynamic assembly of chromosome axes. Dev Cell 33, 94–106 (2015).

39. Shintomi, K. & Hirano, T. A Sister Chromatid Cohesion Assay Using Xenopus Egg Extracts. Methods Mol Biol 1515, 3–21 (2017).

40. Shintomi, K. & Hirano, T. Reconstitution of Mitotic Chromatids In Vitro. Curr Protoc Cell Biol 79, e48 (2018).

